# An evolutionary landscape of sesame: chromosomal variation, allopolyploid speciation and metabolic specialization

**DOI:** 10.64898/2026.03.27.714335

**Authors:** Hiroyuki Tanaka, Eiichiro Ono, Tenta Segawa, Jun Murata, Hiroki Takagi, Yuto Uegaki, Hiromi Toyonaga, Akira Shiraishi, Motoshige Takagi, Atsushi Toyoda, Kyoko Sato, Tatsuya Wakasugi, Manabu Horikawa, Makoto Kawase, Takehiko Itoh, Masayuki P. Yamamoto

**Author notes:** These authors equally contributed to this work (Hiroyuki Tanaka, Eiichiro Ono, Tenta Segawa). These authors are equally responsible for this work (Makoto Kawase, Takehiko Itoh, Masayuki P. Yamamoto).

## Abstract

Sesame (*Sesamum indicum*) is one of the earliest domesticated oilseed crops and is valued for antioxidant lignans that stabilize oil quality. However, the genomic and evolutionary history of the genus *Sesamum*, including the origin of its allotetraploid relative *S. radiatum* and the diversification of lignan metabolism, remains poorly understood owing to limited chromosome-scale genomic resources. Here we present chromosome-level genome assemblies for three wild *Sesamum* species, two *Ceratotheca* species and a Japanese sesame cultivar to reconstruct genome and karyotype evolution across the *Sesamum–Ceratotheca* complex. Comparative analyses show that the derived *x*=16 lineage originated from an ancestral *x*=13 karyotype through chromosome fission, fusion and translocation, whereas another *x*=13 lineage underwent extensive restructuring associated with retrotransposon expansion. Phylogenomics places *Ceratotheca* within the *x*=16 *Sesamum* clade and reveals that *S. radiatum* originated through hybridization involving a *C. sesamoides*-like ancestor. The antioxidative lignan gene *CYP92B14* was reintroduced via the BB progenitor, linking hybridization with restoration of oil-stabilizing metabolism during sesame evolution.

## INTRODUCTION

Understanding how genome evolution, hybridization and specialized metabolism interact remains a central challenge in crop evolution. Sesame (*Sesamum indicum* L.), a member of the Pedaliaceae, is among the earliest domesticated oilseed crops^1–3^, provides an informative system for examining these processes. Originating in the Indian subcontinent^1–4^ and being spread through early transcontinental trade, sesame became a valued commodity across the Indian Ocean and Asian regions^1–3^. Historical sources suggest that rivalry over sesame oil and seed trade shaped regional economies and politics during the medieval period, underscoring its deep economic and cultural significance^3^. Beyond its historical role, sesame is notable for its phenylpropanoid-derived lignans—(+)-sesamin, (+)-sesamolin and (+)-sesaminol—which confer remarkable oxidative stability and health-promoting properties to its seed oil^5,6^. The genus *Sesamum* comprises about 20 species, including the cultivated sesame (*S. indicum)*, the semi-domesticated *S. radiatum*, and other wild taxa distributed mainly across sub-Saharan Africa^1,2,4,7^. Despite decades of morphological and cytogenetic study, phylogenetic relationships within the genus remain poorly resolved. Species have been grouped by basic chromosome number (*x* = 13 or *x* = 16)^8^, with early reports even proposing *x* = 8 for some taxa^9,10^. Persistent inconsistencies in chromosome counts and misapplied names—such as the confusion between *S. indicum* subsp. *malabaricum* and *S. mulayanum*—have hindered clear classification^1,4,11^. The morphological similarity between *S. radiatum* and an accession tentatively identified as *S. schinzianum* No. 124 adds to this ambiguity. The accession has now been re-identified as *S. radiatum* by Bedigian based on morphological characters^12^. In addition, genomic comparisons of these taxa in the present study reveal minimal differentiation between them. The genera *Sesamum* and *Ceratotheca* are considered the closest relatives but distinct lineages^1,7^. Chloroplast data have thus offered a partial framework^13^; however, organellar markers alone are insufficient to resolve the nuclear genomic relationships and past hybridization events within this complex. The first whole-genome sequence of the *S. indicum* ‘Zhongzhi No. 13’ was reported^14^, followed by genomes of several cultivated sesame varieties^15,16^. Recently, the genome sequence of *S. schinzianum* ‘Gangguo’ with 2*n* = 64 chromosomes suggested an allotetraploid origin, possibly resulting from hybridization between two species in the *x* = 16 group^17^. However, for the reasons stated above and the data described below, we consider this accession should be classified as *S. radiatum* not *S. schinzianum*. Miao et al. (2024)^8^ revealed the whole-genome sequences of six wild sesame species (*S. alatum, S. angustifolium, S. angolense, S. calycinum, S. latifolium,* and *S. radiatum*). They classified the diploid species into four groups: AA genome (*S. latifolium*: 2*n* = 2*x* = 32), BB genome (*S. angustifolium, S. angolense, S. calycinum*: 2*n* = 2*x* = 32), CC genome (*S. indicum*: 2*n* = 2*x* = 26), and DD genome (*S. alatum*: 2*n* = 2*x* = 26), and also suggested that *S. radiatum* is an allotetraploid (2*n* = 4*x* = 64) with two diploid genomes (AABB), and the species originated from hybridization between *S. latifolium* (AA) and *S. angustifolium* (BB), followed by chromosome duplication. At the pathway level, lignan biosynthesis is well characterized: CYP81Q1 catalyzes (+)-sesamin formation^18^, and CYP92B14 drives its oxidative conversion to (+)-sesamolin and (+)-sesaminol^19^. (+)-Sesamin is the predominant lignan in wild relatives^20^, while the oxidized sesamin derivatives that attribute to antioxidative property in seed oil^6^ are prominent in *S. indicum* and *S. radiatum*^21^. The highly diverged specialized lignans, (+)-alatumin and (+)-2-episesalatin are reported in *S. alatum* (DD)^22,23^. Yet the genomic architecture underlying this pathway—its copy-number dynamics, chromosomal context and lineage-specific evolution—remains unresolved. Here we present chromosome-level genome assemblies for three wild *Sesamum* species (*S. alatum, S. angustifolium* and *S. latifolium*), two *Ceratotheca* species (*C. triloba* and *C. sesamoides* also known as false sesame), and a Japanese cultivar of *S. indicum* (‘Masekin’). Together, these chromosome-scale assemblies refine nuclear phylogenetic relationships within the *Sesamum–Ceratotheca* complex, elucidate the hybrid origin of *S. radiatum* between *C. sesamoides* and *S. angustifolium*, and reveal genomic signatures underlying the diversification of lignan biosynthesis.

## RESULT

### Chromosome number variation in the genera *Sesamum* and *Ceratotheca*

We determined the somatic metaphase chromosome numbers of *Sesamum* and *Ceratotheca* species whose genomes were sequenced in this study, and performed karyotype analyses for a subset of these species (Fig. 1, Supplementary Table 1 and Supplementary Fig. 1). As previously reported, *S. alatum* and *S. indicum* both have 2*n* = 26, while *S. angustifolium* and *S. latifolium* have 2*n* = 32, and *S. radiatum* has 2*n* = 64^4,8–10,13^. The two *Ceratotheca* species have 2*n* = 32^9,10,13^. *Sesamum* and *Ceratotheca* species are classified into two groups based on their basic chromosome numbers: *x* = 13 and *x* = 16. Thus, species with 2*n* = 26 are considered diploids with *x* = 13 (2*n* = 2*x* =26), while those with 2*n* = 32 and 64 are classified as diploids (2*n* = 2*x* = 32) and tetraploids (2*n* = 4*x* = 64), respectively.

**Fig. 1.**
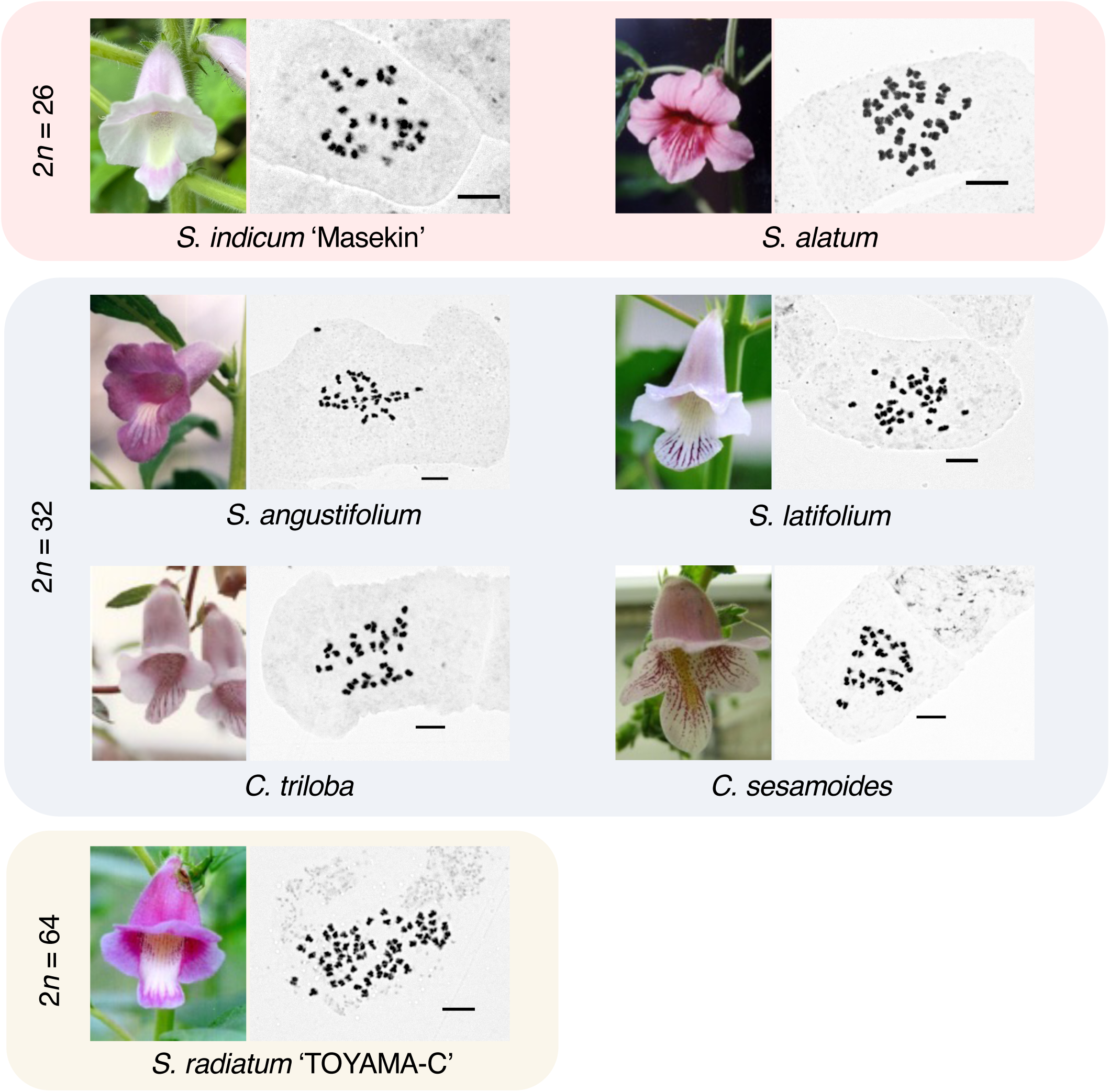
Flower morphology and chromosome observations of the sesame species analyzed in this study. The scale bars in the chromosome observation images represent 5 μm.

### Genome sequences and phylogenetics

We generated de novo assemblies for six species (Supplementary Table 2). PacBio long-read sequencing (PacBio CLR) and Hi-C technologies produced high-quality chromosome-scale genome assemblies for *S. alatum*, *S. angustifolium*, and *C. sesamoides*. These assemblies contained sequences corresponding to 13 chromosomes in *S. alatum*, 16 chromosomes in *S. angustifolium* and *C. sesamoides*, matching the observed karyotypes (Figs. 1 and 2A, Supplementary Fig. 1 and Table 3). Moreover, genome assemblies were also obtained for *S. indicum* ‘Masekin’, *S. latifolium,* and *C. triloba*. Estimated assembly sizes were consistent with *k*-mer profiles, and Benchmarking Universal Single-Copy Orthologs (BUSCO) completeness exceeded 98% in all cases (Supplementary Table 3), indicating high completeness of the assemblies and a quality comparable to or better than those reported in previous studies^8^. Of note, genome size did not correlate with chromosome number; *S. alatum* possesses the largest genome (ca. 537 Mb) among our samples, largely due to extensive expansion of Angela-type LTR/Copia retrotransposons (Supplementary Fig. 2, Supplementary Texts 1 and 2)

**Fig. 2.**
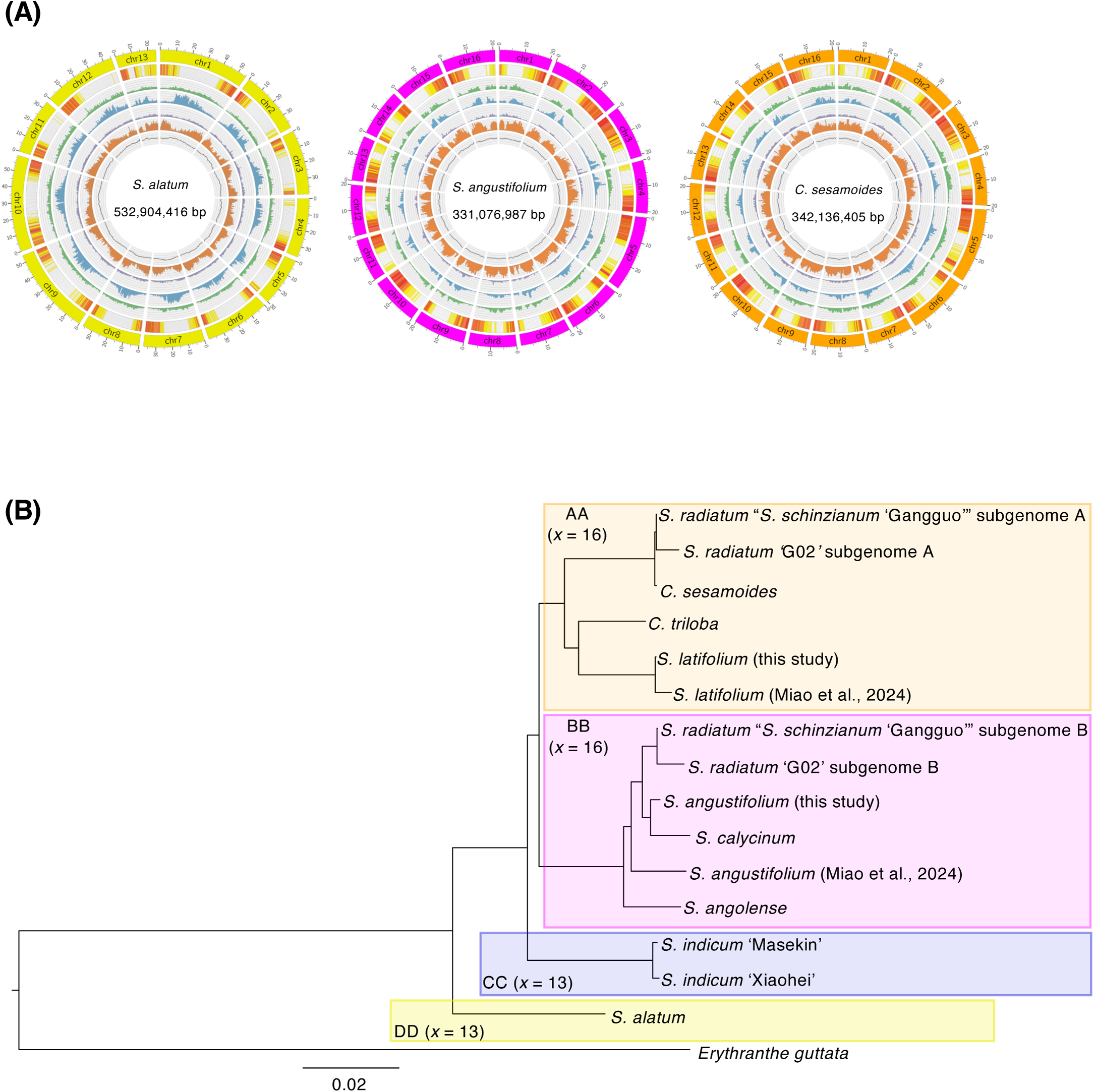
The phylogenetic relationships of *Sesamum* and *Ceratotheca* species. (**A**) Genomic landscapes of *S. alatum*, *S. angustifolium*, and *C. sesamoides* assemblies. From outer to inner rings, tracks represent pseudochromosomes, gene density (1–250 per 500 kb), LTR/Gypsy element density (0–250 per 500 kb), LTR/Copia element density (0–250 per 500 kb), other LTR element density (0–250 per 500 kb), non-LTR element density (0–250 per 500 kb), and GC content (10%–60% per 500 kb). (**B**) Phylogeny of *Sesamum* and *Ceratotheca* species inferred from 178 loci from BUSCO single-copy orthologs conserved across all species and sharing an identical exon number were used. *E. guttata* was used as an outgroup. The scale bar indicates the nucleotide substitutions per site.

A maximum-likelihood phylogeny analysis based on single-copy BUSCO genes placed the *x* = 16 clade as derived relative to *x* = 13, with *S. alatum* (DD) sister to *S. indicum* (CC). Within *x* = 16, two sublineages were recovered: BB (*S. angustifolium*) and AA (*S. latifolium, C. triloba, C. sesamoides*) (Fig. 2B). The subgenome A of the previously published *S. radiatum* “*S. schinzianum* ‘Gangguo’” grouped with *C. sesamoides* rather than *S. latifolium.* Genome-wide dot plots corroborated near one-to-one collinearity between the *S. radiatum* subgenome A and *C. sesamoides*, whereas similarity to *S. latifolium* was reduced and fragmented collinearity. The B subgenome aligned with *S. angustifolium* as previously reported^8^ (Fig. 2B and Supplementary Fig. 3). These results were consistent with the BUSCO-based phylogenetic topology (Fig. 2B).

### Genome rearrangements and structural divergence

Phylogenomic analyses together with chromosome counts support *x* = 13 as the ancestral karyotypic state, from which the *x* = 16 lineage subsequently emerged (Fig. 2B). Comparative genome analyses further indicate that overall gene order is broadly conserved between the *x* = 13 and *x* = 16 lineages, although lineage-specific structural differences are evident (Fig. 3 and Supplementary Fig. 4A). In particular, gene synteny is largely conserved between the AA and BB genomes, whereas substantial divergence is observed between the CC and DD genomes (Fig. 3). Notably, the DD genome of *S. alatum* displays the largest rearrangement distances among all ingroup lineages (Supplementary Fig. 4A). Macro-synteny analyses further reveal that chromosomes 14, 15, and 16 of *x* = 16 taxa (*S. angustifolium*; BB and *C. sesamoides*; AA) are composite mosaics, each showing extensive synteny with regions corresponding to chromosomes 1, 2, and 8 of *S. indicum* (CC), respectively (Fig. 3 and Supplementary Fig. 5). These syntenic relationships indicate that the three additional chromosomes in *x* = 16 taxa cannot be explained by simple chromosomal fissions from *x* = 13 ancestors, but instead reflect multiple chromosomal fission and fusion events. Consistent with these observations, a rearrangement-based phylogeny recapitulates the BUSCO-supported ortholog-based topology (Fig. 2B and Supplementary Fig. 4B), supporting a scenario in which structural divergence between *x*-groups contributed to the hybrid sterility reported in inter-group crosses^4,10^, thereby potentially promoting reproductive isolation during the evolution of sesame.

**Fig. 3.**
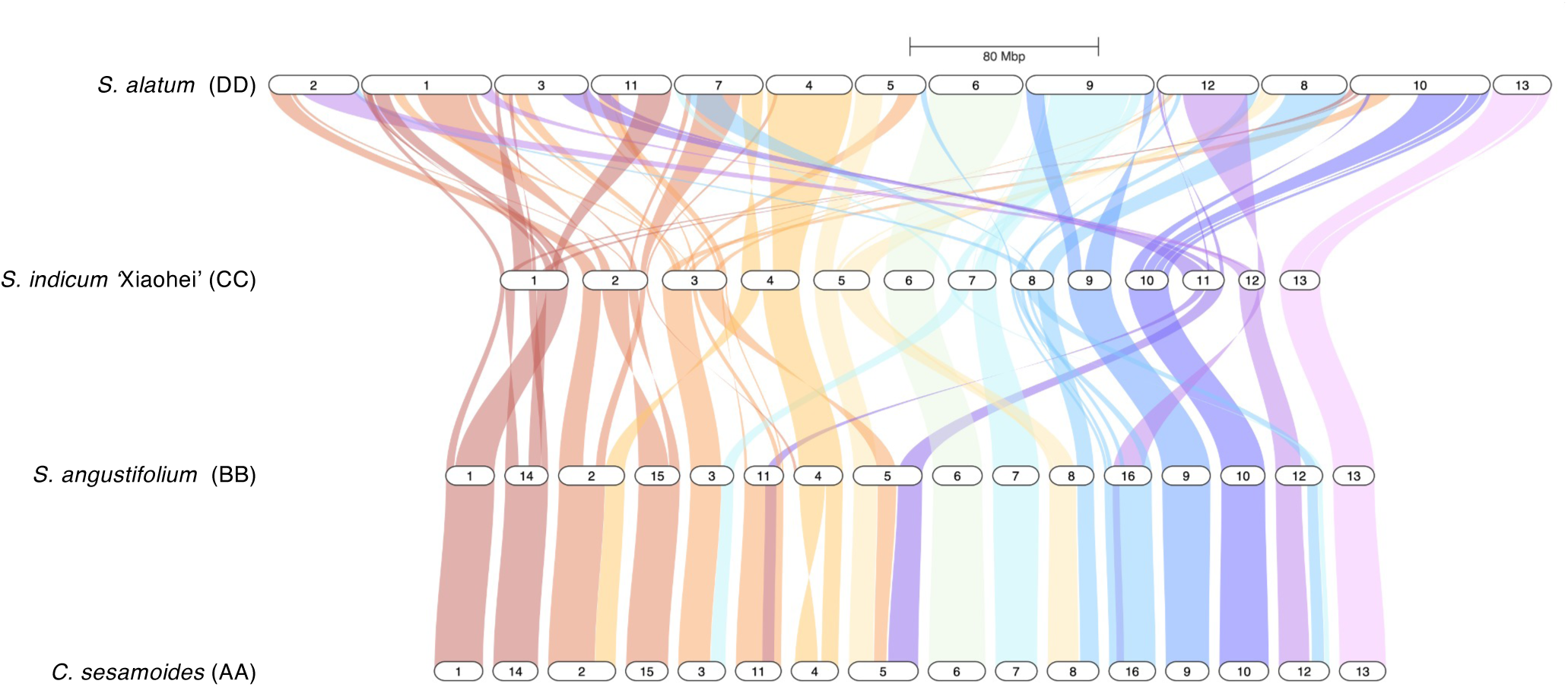
Comparative chromosome structure across *Sesamum* and *Ceratotheca* species. Chromosomal collinearity for four representative species (*S. alatum*, *S. indicum* ‘Xiaohei’, *S. angustifolium*, *C. sesamoides*) is visualised with Sankey ribbons. Ribbon colour corresponds to the originating chromosome in *S. indicum* ‘Xiaohei’; uniform colours denote direct orthology, whereas mixed colours reveal inter-chromosomal exchanges. Chromosome numbers appear in white blocks.

### Allopolyploid speciation of *S. radiatum*

To test whether the *S. radiatum* “*S. schinzianum* ‘Gangguo’” genome is distinct from *S. radiatum*, we remapped short reads from nine *S. radiatum* accessions (including one previously labeled *S. schinzianum* ‘TOYAMA-W’) to that assembly. Beyond balanced mapping to A and B subgenomes, genome-wide SNP concordance shows that *S. radiatum* “*S. schinzianum* ‘Gangguo’” and ‘TOYAMA-W’ definitely falls within *S. radiatum* variation rather than forming a separate clade (Supplementary Fig. 6), indicating conspecificity (Fig. 4A). Reads from candidate diploids mapped preferentially to one subgenome: *S. angustifolium* to subgenome B and *C. sesamoides* to subgenome A; *S. latifolium* showed lower, mixed mapping (Fig. 4A). Together with ortholog phylogenies, synteny and TE composition, these results support an AABB origin from ancestors closest to *C. sesamoides* (AA) and *S. angustifolium* (BB) (Fig. 2B and Supplementary Figs. 2, 3, 6).

**Fig. 4.**
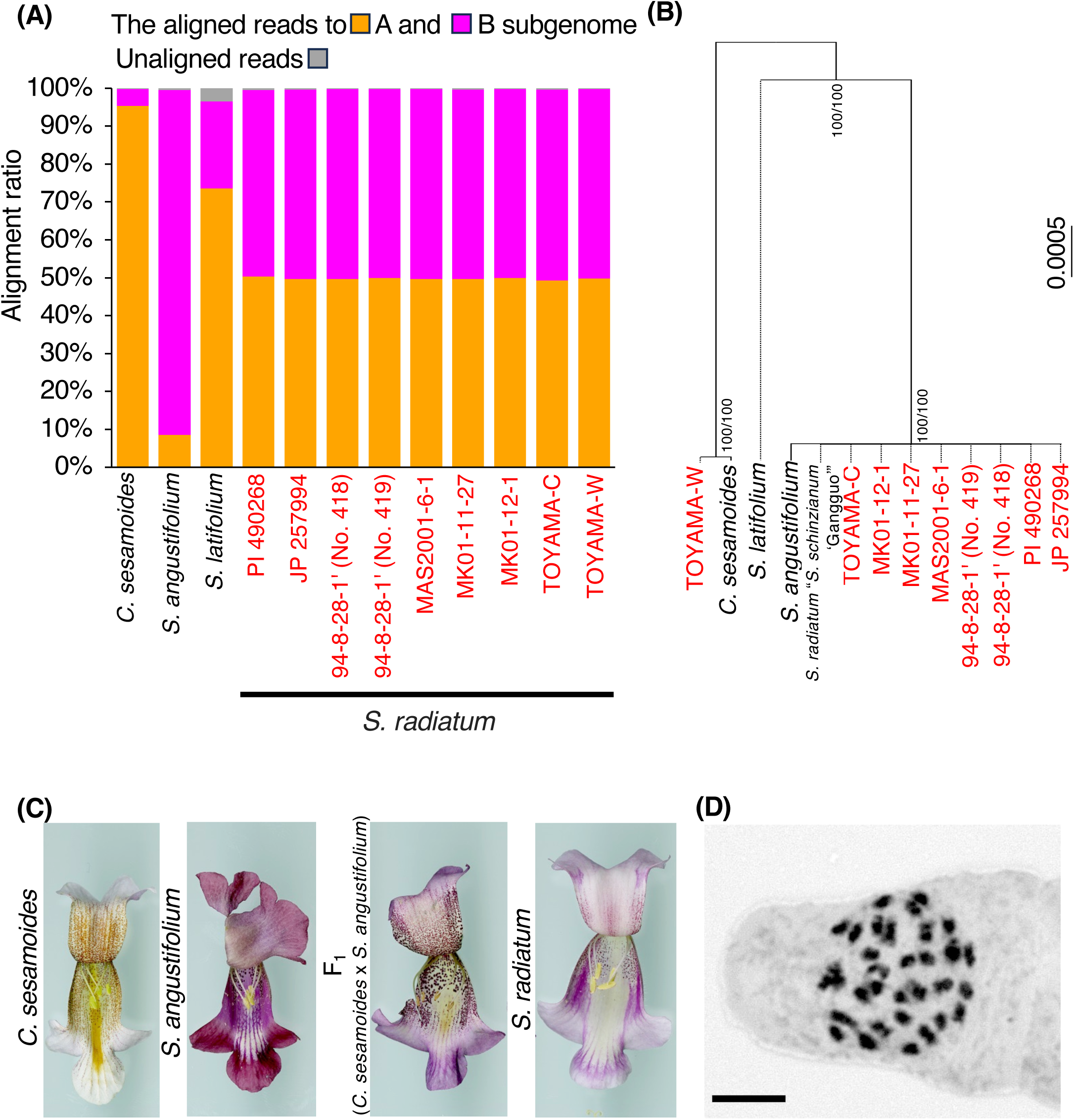
Allopolyploid origin of *S*. *radiatum*. (**A**) The ratio of Illumina short reads aligned to the A and B subgenomes of *S. radiatum* “*S*. *schinzianum* ‘Gangguo’” thorough resequencing. (**B**) Phylogenetic tree based on SNP positions in the chloroplast genome. The line name of *S*. *radiatum* is highlighted in red. Numbers on the nodes indicate bootstrap value from 1,000 replications (left), and SH-aLRT value (right). The scale bar indicates the nucleotide substitutions per SNP site. (**C**) Flower phenotype of *C*. *sesamoides*, *S*. *angustifolium*, their F_1_ and *S. radiatum*. Flowers were opened longitudinally from the tip to the base. (**D**) The chromosome observation of F_1_ plant (*C. sesamoides* x *S. angustifolium*). The scale bar = 5 μm.

Organelle genomes of *S. radiatum* accessions comprised two types: most matched *S. angustifolium*, whereas accession TOYAMA-W matched *C. sesamoides* (Fig. 4B and Supplementary Fig. 7). This is consistent with reciprocal crosses during formation. Divergence-time estimates from fourfold-degenerate sites (BEAST) place the *C. sesamoides*–subgenome A split on the order of 10⁴ years (0.0008-0.0467 MYA) and the *S. angustifolium*–subgenome B split on the order approximately 10⁵ years, within 0.18-0.39 MYA (Supplementary Fig. 8). Such shallow timescales covering the period of human activity support a recent (and potentially recurrent) allopolyploid origin. Moreover, divergence between the AA and BB genomes of *Sesamum* and *Ceratotheca* species was estimated to be within 3.5-4.1 MYA, while the divergence of the *x* = 13 and *x* = 16 groups was within 3.8-4.5 MYA. Therefore, divergence of the A and B genomes occurred shortly after the emergence of species with 2*n* = 2*x* = 32 chromosomes.

### Experimental crosses between *C. sesamoides* and *S. angustifolium*

Reciprocal *S. angustifolium* (BB) × *C. sesamoides* (AA) crosses produced viable F₁ seeds. *CYP81Q* genotyping confirmed parental heterozygosity, and plastid markers verified both maternal directions (Fig. 4C and Supplementary Fig. 9). F₁ plants exhibited intermediate leaf and flower morphology but were almost sterile, however, a small number of capsules were observed to develop during the later stages of growth. We obtained a small number of seeds from F_1_ plant, upon crossing with *S. radiatum* pollen, indicating that they were fertile at least as the seed parent. The chromosome number of 2*n* = 32 (Fig. 4D) suggests that genome doubling would be required to restore fertility, consistent with the allotetraploid *S. radiatum* (2*n* = 64). Ability of the reciprocal crosses was in accordance with the presence of the two organelle types in natural variety of *S. radiatum* (Fig. 4B and Supplementary Fig. 7).

Seed comparisons showed that *S. radiatum* has larger, heavier seeds than *S. angustifolium* and is similar to *C. sesamoides* and *S. indicum* (Supplementary Fig. 10). Distribution records show broad overlap between *S. angustifolium* and *C. sesamoides* across Central/East Africa, providing possible opportunities for hybridization, whereas *S. latifolium* is more geographically restricted (Supplementary Fig. 11). Together with the phylogenomic evidence herein, these data point to *C. sesamoides* rather than *S. latifolium* as the more plausible ancestor of an allotetraploid sesame, *S. radiatum*.

### Metabolic specialization in oxidized sesamin

Lignan profiling detected (+)-sesamin in all diploids except *S. alatum*; oxidized derivatives—(+)-sesamolin, (+)-sesaminol and (+)-sesangolin—were restricted to *S. indicum*, *S. angustifolium* and *S. radiatum* (Fig. 5A and Supplementary Table 5). All wild relatives carried a single CYP81Q ortholog, whereas *S. radiatum* retained two homeologues (SrCYP81Q2 from AA; SrCYP81Q4 from BB), and all tested CYP81Qs except CYP81Q3^23^ were active (+)-sesamin synthases (Supplementary Fig. 12). By contrast, (+)-sesamin-oxidising CYP92B14^19,21^ was present only in CC and BB genomes and in *S. radiatum* (Fig. 5 and Supplementary Figs. 13, 14). Gene synteny comparison showed CYP92B14 is located within a subtelomeric CYP cluster in these genomes and absent from AA and DD lineages. A possible scenario consistent with these data is that CYP92B14 was present on chr8 of the *x* = 13 ancestor, lost from chr16 in the AA lineage after divergence from BB within *x* = 16 (Fig. 5B), and restored in *S. radiatum* via BB-derived hybridization. The subtelomeric context aligns with regions prone to copy-number dynamics, matching the observed lineage-specific pattern.

**Fig. 5.**
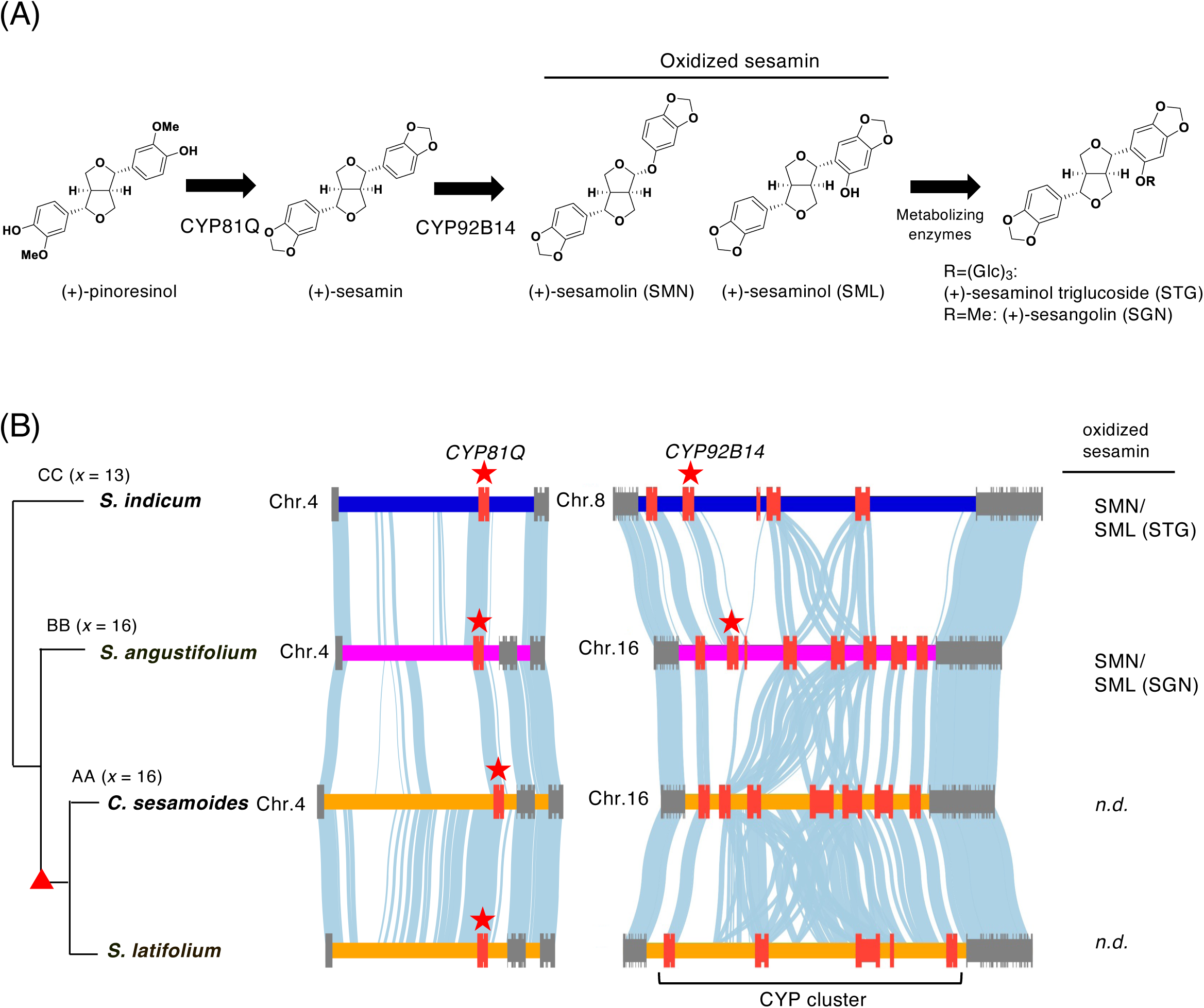
Evolutionary landscape of lignan metabolism genes in *Sesamum* and *Ceratotheca* species. (**A**) Representative biosynthetic pathway of sesame lignans. (+)-Pinoresinol is first converted to (+)-sesamin by a CYP enzyme CYP81Q1. A second CYP, CYP92B14, then oxidizes (+)-sesamin to two major derivatives: (+)-sesamolin (SMN) and (+)-sesaminol (SML). Subsequently, SML undergoes either glucosylation or methylation to form (+)-sesaminol triglucoside (STG) or (+)-sesangolin (SGN), respectively. (**B**) Conservation of the *CYP81Q* and *CYP92B* loci. The cladogram (left) shows the relationships and chromosome numbers of four *Sesamum* species. For each species, colored bars represent the chromosomal segments that harbor the *CYP81Q* locus (chromosome 4) and the *CYP92B14* locus (chromosome 8 in *S. indicum*; chromosome 16 in the 16-chromosome taxa). Red boxes mark the focal CYP genes; black boxes denote other CYP genes; grey boxes indicate neighboring genes. Light-blue ribbons link collinear gene pairs, illustrating conserved micro-synteny. A tandem CYP cluster containing multiple *CYP92B14* paralogues is present on chromosome 16 of the 16-chromosome genomes. The right-hand column summarizes the oxidized sesamin products detected in each species (SMN/SML (STG), SML (SGN), or n.d. = not detected). The red triangle indicates a point of the *CYP92B14* gene loss.

## Discussion

The findings presented here establish a unified framework for understanding how chromosome evolution, hybridization and specialized metabolism interact during crop evolution. By integrating chromosome-scale comparative genomics with experimental hybridization, we resolve the origin of the allotetraploid sesame *S. radiatum* and demonstrate that hybridization can restore a specialized metabolic function that was lost in one parental lineage. This result expands prevailing views of hybridization as a driver of novelty by showing that it can also act as a mechanism for metabolic recovery, with direct consequences for agronomically important traits such as oil stability. More broadly, our study illustrates how structural genome evolution and allopolyploid formation can shape both reproductive isolation and chemical diversity in plants, providing insights that extend beyond sesame to other crop and non-crop systems. By assembling chromosome-scale genomes across *Sesamum* and *Ceratotheca*, we provide an integrated view of genome evolution within the Pedaliaceae. Our analyses show how chromosomal reorganization, retrotransposon activity, and interspecific hybridization together shaped the diversification of *Sesamum* species. Consistent with classical cytogenetic observations^9˒10^ and recent genomic work^8^, the *x* = 16 lineage diverged from an ancestral *x* = 13 group that includes *S. indicum*. The massive change in chromosome numbers was driven not by genome duplication but by multiple chromosomal fission, reciprocal translocations, and pericentromeric rearrangements^24–26^. The chromosomal rearrangements associated with the transition from *x* = 13 to *x* = 16 likely contributed to the hybrid sterility reported in inter-group crosses^4, 10^, thereby accelerating speciation of sesame by reproductive barriers. Chromosome-level genome assemblies from additional *Sesamum* and its related species will be essential to resolve mechanisms of chromosome number variation within this family.

The large, repeat-rich genome of *S. alatum* exemplifies lineage-specific expansion. Its inflated intergenic regions indicate that retrotransposon bursts may have altered local chromatin structure around metabolic gene clusters. Because *S. alatum* also exhibits unique morphology, narrow leaves, long floral tube, winged seeds and specialized lignan chemotypes^4,22,23^, the link between genome inflation and metabolic specialization may be mechanistic rather than coincidental. Together with the contrast between CC and DD genomes within the *x* = 13 group, these observations demonstrate that extensive genome restructuring can accumulate independently of changes in chromosome number. This decoupling underscores that chromosome number evolution and large-scale structural divergence represent separable modes of genome evolution in *Sesamum*.

Our phylogenomic framework also clarifies the long-debated relationship between *Sesamum* and *Ceratotheca*. Morphological analyses^1,7^ and chloroplast phylogenies^13^ have long implied close relationship between the two genera, yet organellar data alone could not resolve their evolutionary divergence. Nuclear phylogenies and genome-wide synteny now place both *Ceratotheca* species firmly within the *x* = 16 *Sesamum* lineage, indicating that the genera form a continuous evolutionary assemblage rather than distinct lineages. This finding supports earlier taxonomic proposals that *Ceratotheca* should be classified within *Sesamum*^13^ and underscores how incomplete lineage sorting and historical introgression can decouple chloroplast and nuclear signals. Most notably, our results resolve the origin of the allotetraploid *S. radiatum* (2*n* = 4*x* = 64). We show that its subgenomes A and B derive from ancestors related to *C. sesamoides* (AA genome) and *S. angustifolium* (BB genome), respectively—contradicting earlier hypotheses assigning the subgenome A to *S. latifolium*^8^. Multiple independent lines of evidence—including whole-genome alignments, ortholog phylogenies, synteny conservation, organellar genotypes, and transposon composition—support *C. sesamoides* as the true parental progenitor. This is consistent with previous reports showing that *S. radiatum* morphologically resembles *Ceratotheca*, indicating a close relationship with the genus^4^. The extremely low divergence between the *C. sesamoides* AA genome and the *S. radiatum* subgenome A (0.0008–0.05 Mya) indicates that polyploidization occurred recently and possibly recurrently within the period of human activity. Population genomic analysis of local *S. radiatum* varieties will clarify this recurrent issue in future research.

Reciprocal crosses between *S. angustifolium* and *C. sesamoides* produced viable but sterile F₁ hybrids resembling *S. radiatum* in morphology (Fig. 4C,D and Supplementary Fig. 9C), showing that genome doubling was required to restore fertility—a canonical route to allopolyploid speciation^27,28^. Given that both parental species are cultivated sympatrically as leafy vegetables across sub-Saharan Africa (Supplementary Fig. 13)^4,7,29–31^, small-scale cultivation and anthropogenic disturbance may have facilitated recurrent hybrid formation, ultimately contributing to the emergence of *S. radiatum* as a seed oil crop with large seed, comparable to that of *S. indicum* (Supplementary Fig. 10).Beyond cytogenetics, our data illuminate the genomic basis of metabolic specialization in *Sesamum*. The CYP enzyme CYP92B14 catalyzes the oxidative rearrangement of (+)-sesamin into oxidized derivatives such as (+)-sesamolin and (+)-sesaminol^19,21^, compounds that confer oxidative stability to sesame oil^6^. We found that in contrast to the highly conserved CYP81Q gene in *Sesamum* and its related species in Lamiales including *Phryma leptostachya* and *Paulownia tomentosa* (Supplementary Fig. 13A), CYP92B14 gene is restricted to the BB and CC genomes and absent from AA and DD lineages (Fig. 5 and Supplementary Figs. 13B, 14), indicating a lineage-specific metabolic specialization from the primary sesame lignan, (+)-sesamin. Comparative genomics reveal the CYP92B14 cluster in subtelomeric regions at approximately 36.0 kb and 29.9 kb from the respective chromosome ends —genomic environments prone to copy-number variation and rapid divergence^32,33^. In *S. radiatum*, this cluster was inherited intact from its BB progenitor *S. angustifolium*, providing a metabolic route by which hybridization transferred the biochemical capacity for (+)-sesamin oxidation. This metabolic specialization would be contributing to the seed oil properties of *S. radiatum*.

Together, these findings reveal a dynamic evolutionary trajectory in which chromosomal rearrangements, transposable element activity, and interspecific hybridization intersect to generate genetic diversity and biochemical novelty in sesame. The nuclear phylogenomic data herein might call for a re-classification of *Ceratotheca* genus in Pedaliaceae family as proposed by chloroplast-based phylogeny^13^. Comparative genomics approaches herein provided the structural basis for diversification and hybridization merged divergent gene pools, enabling new metabolic repertoires. Future integration of population genomics, transcriptomics, and metabolomics will further refine this framework.

## Materials and Methods

### Plant materials

Five species of *Sesamum* (*S. indicum* ‘Masekin’, *S. alatum*, *S. angustifolium*, *S. latifolium*, and nine accessions of *S. radiatum*) and two species of *Ceratotheca* (*C. sesamoides* and *C. triloba*) are used in this study (Fig. 1 and Supplementary Table 1). Among the accessions of *S. radiatum* ‘TOYAMA-W’ was previously classified as *S. schinzianum*; however, it is now correctly identified as *S. radiatum* based on morphological and genomic evidence. The USDA gene bank initially classified *S. angustifolium* (PI 367899) as *S. radiatum*. However, based on a comprehensive analysis of plant morphology, chromosome numbers, and genomic data, we have determined that the correct classification is *S. angustifolium*.

### Chromosome observation

Plants used in this study were grown from stock seeds in an experimental garden and greenhouse at University of Toyama. For chromosome counts, root tips were pretreated with 1.6 mM 8-hydroxyquinoline solution at room temperature for 1 to 1.5 h and kept at 5°C for 15 h. After the appropriate pre-treatment, the root tips were fixed in a mixture of glacial acetic acid and absolute ethyl alcohol (1:3) at room temperature (ca. 25℃) for 1 h, macerated in 1M hydrochloric acid at 60℃ for 10 min, and then washed in tap water. They were stained and squashed in 1.5% lacto-propionic orcein. Fully spread metaphase chromosomes were then observed under microscope.

### Genome and RNA-sequencing

Genomic DNAs for genome sequencing were extracted from mature leaves of each plant. The leaves were frozen and ground in liquid nitrogen. DNA from *S. indicum* ‘Masekin’ were prepared DNeasy Plant Kits (Qiagen) and DNA from *S. latifolium* were isolated according to previous works^19^. DNAs from other species were isolated using the MagExtractor–Plant Genome kit (TOYOBO, Japan) following the manufacturer’s instructions. Total RNAs were extracted from *Sesamum and Ceratotheca* according to previous studies^18,19,34^. Genomic sequences of S*. alatum*, *S. angustifolium*, *C. sesamoides* and *C. triloba* were determined with PacBio single-molecule real-time sequencing and Illumina paired-end short reads. For PacBio sequencing, a Continuous Long Read (CLR) SMRTbell library was constructed with the SMRTbell Express Template Kit 2.0 (Pacific Biosciences, CA, USA) according to the manufacturer’s protocol. The CLR library was size-selected using the BluePippin system (Sage Science, MA, USA) with a lower cutoff of 30 kb. One SMRT Cell 8M was sequenced on the PacBio Sequel II system with Binding Kit 2.0 and Sequencing Kit 2.0. In addition, Illumina paired-end libraries were constructed from fragmented genomic DNA using PCR-free library preparation methods and sequenced on an Illumina NovaSeq 6000, generating 150 bp paired-end reads. Genomic sequences of *S. indicum* ‘Masekin’ and *S. latifolium* were determined with Oxford Nanopore Technologies and Illumina paired-end short reads. For ONT sequencing, high-molecular-weight DNA was prepared and ligation-based sequencing libraries were constructed following the manufacturer’s protocols. Libraries were sequenced on a PromethION. Basecalling was performed using Guppy with default parameters. In addition, Illumina paired-end libraries were prepared from fragmented genomic DNA and sequenced on an Illumina HiSeq X Ten, generating 150 bp paired-end reads. Omni-C libraries (Dovetail Omni-C, Dovetail Genomics, CA, USA) were prepared for *S. alatum*, *S. angustifolium* and *C. sesamoides* from young leaves according to the manufacturer’s instructions and sequenced on an Illumina NovaSeq 6000. The proximity-ligation data were used for Hi-C scaffolding to produce chromosome-scale assemblies.

For RNA-seq, total RNA was extracted from five tissues (leaves, stems, flowers, seed pod, and roots), and strand-specific RNA-seq libraries were constructed following the manufacturer’s protocol. Sequencing was performed on an Illumina NovaSeq 6000, generating 150 bp paired-end reads. The RNA-seq data were used to support gene prediction.

Raw reads for all accessions are deposited under BioProject PRJDB20216; individual read accessions and metadata are listed in Supplementary Table 6.

#### Genome size estimation

The genome size of three *Sesamum* and two *Ceratotheca* species was estimated from Illumina sequencing data using the k-mer-based method. Illumina sequenced reads were filtered using platanus_trim v1.0.7 (http://platanus.bio.titech.ac.jp/pltanus_trim) with default parameters. To remove highly copied reads derived from the chloroplast genome, filtered reads were mapped to the species-specific chloroplast genome assembly generated separately for each taxon, and unmapped reads were extracted. Using the unmapped reads, Jellyfish v2.2.10^35^ was first applied to extract and count canonical k-mers at k = 32. Subsequently, GenomeScope 2.0^36^ was used to estimate haploid genome size and heterozygosity from k-mer count data with parameters of “-k 32”.

### Genome assembly

#### Chloroplast

Illumina-sequenced reads were filtered using platanus_trim v1.0.7 (http://platanus.bio.titech.ac.jp/pltanus_trim) with the following parameters “-q 0.” The trimmed reads were assembled using NOVOPlasty v4.3.1^37^ with the following parameters: “Type = chloro, K-mer = 36, Read Length = 150, Insert size = 550, Platform = Illumina, Single/Paired = PE, Insert size auto = yes”. The chloroplast genome of *S. indicum* was used as the reference10, and psbA chloroplast gene sequences were used as seeds to assemble the plastome.

#### Mitochondrion

We performed a de novo assembly of the trimmed reads using NOVOPlasty v 4.3.1 with the following parameters: “Type = mito_plant, k-mer = 36, Read Length = 250, Insert size = 600, Platform = illumina, Single/Paired = PE, Insert size auto = yes”. The cox1 gene was used fas the seed sequence for assembly. Preliminary draft mitochondrial contigs were successfully generated.

#### Nuclear genome

PacBio sequenced reads were used for genome assembly by the Canu v2.1.1^38^ with parameters of “corOutCoverage=200”. The draft assembly contigs were polished with one rounds of Arrow (Pacific Biosciences) and three rounds of Pilon v1.2.4^39^ with parameters of “”. Nanopore sequenced reads were used for genome assembly by the NextDenovo v2.5.0^40^ with default parameters. The draft assembly contigs were polished with three rounds of NextPolish v1.4.0^41^ using Illumina sequenced reads and Nanopore sequenced reads with default parameters. Haplotypic duplication and redundant contigs were removed from the polished assembly with purge_dups v1.2.5^42^. Organelle contigs were identified by alignment against the already generated mitochondrial and chloroplast genomes using nucmer v4.0.0beta2^43^, and then organelle contigs, redundant contigs and contigs of aberrant GC content (<1 % or >99 %) were removed. To obtain a chromosome assemblies of *S. alatum, S. angustifolium, and C. sesamoides,* we performed Hi-C scaffolding using Hi-C dataset (Omni-C). Bridge sequences were trimmed from the Omni-C reads using cutadapt v4.1^44^. Using the cleaned Omni-C reads, the primary assembly was scaffolded with yahs v1.2a^45^. Briefly, the cleaned Omni-C reads were aligned to the assembly with bwa mem v0.7.17-r118^46,47^. From the mapped reads, valid ligation events were recorded, and PCR duplicates were removed using pairtools v0.3.0 (https://github.com/open2c/pairtools), producing the “.bam” file that was used for scaffolding. We visualized the Hi-C contact map and performed extensive manual curation using Juicebox v1.11.08^48^ to fix mis-assemblies and mis-scaffoldings.

For *S. alatum*, chromosomes were assigned and oriented by whole-genome alignments to S. indicum using MUMmer4 nucmer v4.0.0 (-l 20 -c 65). For each scaffold/chromosome, the counterpart was the *S. indicum* ‘Xiaohei’ chromosome with the largest total aligned length (sum of non-overlapping matches). Strand (+/–) was set by the longer cumulative alignment length in the forward vs reverse orientation. For *C. sesamoides* and *S. angustifolium*, the same procedure was applied against the harmonized *S. radiatum* assembly.

### Gene structural annotation

Gene structural annotation of all assemblies was conducted using a combination of RNA-seq transcriptome sequences for gene structure prediction, homology-based gene prediction using protein sequences of related species and ab initio gene prediction. For the transcriptome-based gene prediction, RNA-seq reads were filtered by Platanus_trim (http://platanus.bio.titech.ac.jp/pltanus_trim) to remove adaptor and low-quality sequences. De novo transcriptome assembly was performed with Trinity v2.8.4^49,50^ and Oases v0.2.8^51^ and redundant sequences were trimmed with CD-HIT v4.6^52^. The assembled sequences were aligned using Gmap v2019-02-1^53^ and complete open reading frames (ORFs) were predicted by TransDecoder v5.0.2 (https://github.com/TransDecoder/TransDecoder). In addition to de novo assembly, gene prediction was also performed by mapping the RNA-seq reads to the genome. The filtered RNA-seq data were mapped to the genome sequences using HISAT2 v2.2.1^54^, and the mapped transcripts were assembled using StringTie v2.1.7^55^. ORF regions were then identified with TransDecoder. For the homology-based method, protein sequences of *Olea europaea* var. sylvestris (NCBI accession No: GCF_002742605.1), *S. indicum* (NCBI accession No: GCF_000512975.1), Salvia splendens (NCBI accession No: GCF_004379255.2), *Erythranthe guttata* (NCBI accession No: GCF_000504015.1), *Solanum lycopersicum* (NCBI accession No: GCF_000188115.4), and *Antirrhinum majus* (Snapdragon Genome Database: http://bioinfo.sibs.ac.cn/Am/index.php) were used. They were aligned to the genome sequences using Spaln v2.3.3^56^ to predict gene structures. For the ab initio-based method, AUGUSTUS v3.3.2^57,58^ and SNAP v2006-07-28^59^ were used. Both programs were trained on gene models derived from 1,000 genes predicted by the transcriptome-based method. Finally, all predicted gene candidates were merged using the GINGER pipelin^60^.

### Phylogenetic Analysis Based on Single-Copy Orthologs

Whole-genome sequences from 11 taxa belonging to the genera *Sesamum* and *Ceratotheca* were analyzed, including *C. sesamoides*, *C. triloba*, *S. angustifolium* (this study), *S. angustifolium*^5^, *S. calycinum*, *S. angolense*, *S. alatum*, *S. latifolium*, and *S. indicum* ‘Masekin’ and ‘Xiaohei’, as well as the A- and B-subgenomes of *S. radiatum*. To root the phylogenetic tree, Erythranthe guttata was included as an outgroup. Single-copy orthologous genes were identified from the whole-genome sequences using BUSCO v5.4.3^61^ in genome mode. From these, only ortholog groups retained in all species and sharing an identical number of exons were selected for phylogenetic analysis. This filtering resulted in a total of 178 orthologous loci. Multiple sequence alignments were performed for each ortholog group using MAFFT v7.407^62^. Alignments were trimmed using trimAl (option:-automated1) to remove positions containing gaps (“−”) or ambiguous characters (“X”). The filtered alignments were concatenated into a single supermatrix, which was used for phylogenetic reconstruction. A maximum-likelihood phylogenetic tree was inferred using RAxML v8.2.12^63^ with the JTT substitution model and a gamma distribution of rate heterogeneity (option:-m PROTGAMMAJTT).

### Rearrangement-based phylogenetic analysis

Whole-genome assemblies for eleven species (*C. sesamoides*, the A- and B-subgenomes of *S. radiatum* “*S. schinzianum* ‘Gangguo’”, *S. angustifolium*, *S. indicum* ‘Baizhima’ and ‘Xiaohei’, *S. alatum*, *Avicennia marina*, *Andrographis paniculata*, *Mentha longifolia*, *Salvia miltiorrhiza*, and *Ballota nigra*) were first screened with BUSCO v5.4.3 in genome mode against the eudicots_odb10 dataset to obtain single-copy orthologs. These orthologs served as anchors for CHROnicle v1.6: SyChro delineated synteny blocks of at least two collinear genes, merging any overlapping segments, and the resulting breakpoint data were analysed using PhyChro^64^ to infer rearrangement-based phylogenetic relationships. In addition, rearrangement distances between all pairs of genomes were calculated using ReChro, based on the synteny block and breakpoint information derived from SynChro. These pairwise rearrangement distances were used to generate distance matrices and heatmaps, enabling quantitative comparison of chromosomal structural divergence among lineages. The two subgenomes of the allotetraploid *S. radiatum* “*S. schinzianum* ‘Gangguo’” were treated as independent operational taxonomic units.

### Genome-wide dot-plot analysis of orthologous gene pairs

Orthologous gene pairs were identified by reciprocal best-hit BLASTP searches between genomes using predicted protein-coding sequences. Only reciprocal best-hit pairs were retained for downstream analyses. For each genome, genes were ordered along pseudochromosomes according to their genomic coordinates, and cumulative gene indices were assigned sequentially chromosome by chromosome. Genome-wide dot plots were generated by plotting cumulative gene indices of orthologous gene pairs between query and reference genomes. Each dot corresponds to one orthologous gene pair and is colored according to BLASTP percent identity. Chromosome boundaries were indicated by dotted grid lines to delineate transitions between pseudochromosomes. All dot plots were rendered using custom scripts to ensure consistent scaling and visualization across species comparisons.

### Divergence-Time Estimation

We identified 99 single-copy genes using OrthoFinder version 2.5.5^65^ and aligned their amino acid sequences with Clustal Omega version 1.2.4^66^. The amino acid sequences were then converted to nucleotide sequences using PAL2NAL version 14.1^67^. From these, 5743 biallelic SNPs were selected at 4-fold degenerate sites for divergence time estimation with BEAST version 2.7.5^68^. BEAUti in BEAST was configured with the following parameters: Substitution Rate (estimate)=’Tree’, Gamma Category Count=4, Shape (estimate)=‘True’, Substitution Model=’JC69’, Clock Model=’Strict Clock’, and MCMC Chain Length=10,000,000, with a calibration point. The calibration times were referenced from TimeTree and set as follows: *Arabidopsis* and *Erythranthe* at 111.4-123.9 Mya, *Erythranthe* and *Sesamum* at 32.6-63 Mya.

Genomic evaluation of *S. radiatum* “*S. schinzianum* ‘Gangguo’”

Illumina short reads from *C. sesamoides*, *S. angustifolium,* and nine other accessions of *S. radiatum* were mapped to the sequenced *S. radiatum* “*S. schinzianum* ‘Gangguo’” assembly using BWA v0.7.17. Alignments were parsed to assign each primary read to the A or B subgenome according to the annotated contig identifiers, and the subgenome-specific alignment ratio was computed as (reads assigned to subgenome)/(total primary mapped reads) using a custom script.

### Genome sequencing, variant calling, and phylogenetic analysis for *S. radiatum*

Paired-end Illumina reads for *Sesamum radiatum* accessions and diploid comparators were adapter- and quality-trimmed using fastp (v0.23.x)^69^ and mapped to the S. radiatum reference genome using BWA-MEM (v0.7.17) with sample-specific read-group tags. The resulting BAM files were processed with SAMtools (v1.20)^70,71^: mate information was corrected with fixmate, alignments were coordinate-sorted, and PCR duplicates were identified and removed with markdup. We retained uniquely mapped primary alignments by excluding unmapped reads, secondary alignments, supplementary alignments, and QC-failed reads (SAMtools flags 0x904) and requiring MAPQ ≥ 1. Because S. radiatum is an allotetraploid, we analyzed the A and B subgenomes (chromosomes chr1A–chr16A and chr1B–chr16B) separately to avoid confounding variation between homeologous chromosomes. Variants were called per sample using GATK HaplotypeCaller (v4.6.2.0) in GVCF mode assuming disomic inheritance (ploidy = 2) within each subgenome, then jointly genotyped across samples with GenotypeGVCFs^72,73^. Raw variant calls were filtered to retain high-confidence biallelic SNPs: we retained SNP-type variants only, set low-quality genotypes (DP < 5 or GQ < 10) to missing, applied site-level quality filters (QUAL ≥ 25, MQ ≥ 25, QD ≥ 1.5), removed sites with ≥ 27.3% missing data, retained only biallelic sites, and excluded monomorphic sites (AC = 0 or AC = AN). To eliminate ambiguity arising, we retained only sites where all samples showed homozygous genotypes (0/0 or 1/1), excluding any site containing heterozygous calls. This conservative approach yielded 30,149 SNPs (genome A) and 158,240 SNPs (genome B) for phylogenetic analysis. Maximum-likelihood phylogenies were inferred independently for each subgenome using IQ-TREE 2 (v2.3.5)^74^ under the GTR+G4 substitution model with 1,000 ultrafast bootstrap and 1,000 SH-aLRT replicates. Trees were visualized in FigTree (v1.4.4)^75^.

### Genotyping of F₁ hybrids from an interspecific cross

Genomic DNA was extracted from *S. angustifolium*, *C. sesamoides*, and their F₁ hybrids as described above. PCR amplification was performed using KOD FX neo (TOYOBO) according to the manufacturer’s instructions. The primer sets used were as follows: CYP81F (5′-CGAAAGTTGTGCGATCTTGA-3′) and CYP81R (5′-GAGATTCCTGCAACGAAAGC-3′) for *CYP81Q*, and e (5′-GGTTCAAGTCCCTCTATCCC-3′) and f (5′-ATTTGAACTGGTGACACGAG-3′) for the chloroplast *trnL-F* spacer^76,77^.

PCR products were purified using ExoSAP-IT (Thermo Fisher Scientific) and sequenced with the BigDye Terminator v3.1 Cycle Sequencing Kit (Thermo Fisher Scientific), following the manufacturer’s instructions and then analyzed in a 3500 Genetic Analyzer (Thermo Fisher Scientific). Primers used for Sanger sequencing were 5′-AATGATCAACCATGGTGTTCT-3′ for *CYP81Q* and primer e for the *trnL-F* spacer.

### Lignan profile analyses

The filtered sample solution was analyzed using an ion-trap time-of-flight mass spectrometer (Shimadzu LCMS-IT-TOF, Shimadzu Corp., Kyoto, Japan) equipped with a photodiode array detector (Shimadzu, Japan). Each component was separated using a YMC Triart C18 column (TA12S03-1503WT, 150 mm × 3 mm, 3 µm i.d.) with mobile phases A, 0.1% HCO_2_H-H_2_O; and B, 0.1% HCO2H-CH3OH in a linear gradient elution (0-10-25-35-35.01-40 min and 30-50-90-90-30-30% B, respectively) at a flow rate of 0.3 mL min−1.

### Phylogenetics of sesame CYP81Q-related genes

CYP81Q-related orthologs were amplified with genomic DNA extracted from each plant by PCR with the specific primer set (Supplementary Table 7) and the nucleotide sequences were determined. Sequencing reactions were conducted with BigDye-terminator cycle sequencing kit (Thermo Fisher Scientific). Then, the sequencing reaction mixtures were analyzed in a 3500 Genetic Analyzer (Thermo Fisher Scientific). The unrooted phylogenetic tree of CYP81Q-related genes from sesame-related plants at the amino acid level was constructed using the Neighbor–Joining method with SEAVIEW software (version 4.2.4)^78^. The amino acid sequences were aligned by taking into consideration codon positions using CLUSTAL W^79.^ All positions containing gaps and missing data were eliminated from further analysis. A Neighbor–Joining tree was reconstructed by SEAVIEW using a matrix of evolutionary distances calculated by Poisson correction for the multiple substitutions. The reliability of the reconstructed tree was evaluated by a bootstrap test with 1,000 replicates. The accession numbers of the genes subjected to phylogenetic analysis are as follows: SiCYP81Q1, AB194714; SrCYP81Q2, AB194715; SlatCYP81Q2, LC865683; CtriCYP81Q2, LC865684; CsesCYP81Q2, LC865687; SalaCYP81Q3, AB194716; StriCYP81Q3, LC865685; SrCYP81Q4, LC865682; SangCYP81Q4, LC865686; PtCYP81Q116, LC865681; and PlCYP81Q38, AB923911.

### Bioassay experiments in yeast cells and HPLC analysis

Single colonies of yeast strains expressing the ORF sets SrCYP81Q2+CPR1, SrCYP81Q4+CPR1, CtCYP81Q2+CPR1, or SlCYP81Q2+CPR1 were cultured in 3 ml synthetic defined liquid medium overnight at 30 °C with rotary shaking at 120 r.p.m. 50 µl of the stationary phase culture obtained after overnight incubation was transferred to 1 ml of fresh medium in 24-well plates supplemented with 100 µM of the designated lignan as a substrate in the experiment. The cultures were further incubated for 24 h at 30 °C with rotary shaking at 120 r.p.m. 50 µl of the evenly dispersed mixture of the cells and the medium was collected and mixed with 50 µl of acetonitrile. The samples were centrifuged at 21,000×g for 10 min, and the supernatant was filtered through a Millex-LH (Waters, Japan) syringe filter and subjected to HPLC analysis.

HPLC analysis was performed using a 3.0 × 75 mm i.d., 2.7 um Cortecs C18+ (Waters, Japan) at 40 °C. The injection volume was 1 μl. The mobile phase consisted of solvent A containing 0.1% (v/v) trifluoroacetic acid in distilled water and solvent B containing 0.1% (v/v) trifluoroacetic acid in acetonitrile at a flow rate of 1.25 ml/min. The linear gradient used was as follows: 0−1.4 min, 30−80% B; 1.4−1.8 min, 80% B; 1.8−2 min, 80-30% B; 2-2.5 min, 30%. Peaks were detected by fluorescence detector (Ex. 280 nm, Em. 340 nm). The identity of the reaction products was confirmed by comparing the retention time and parent mass with authentic lignan standards.

### Species distribution data and mapping

Occurrence data for *C. sesamoides*, *S. radiatum*, *S. angustifolium*, and *S. latifolium* were retrieved from the Global Biodiversity Information Facility (GBIF; https://www.gbif.org) on 7 May 2025. For each species, occurrence records containing valid geographic coordinates (latitude and longitude) were downloaded in Darwin Core Archive format. Only records with non-missing values for decimalLatitude and decimalLongitude were retained for analysis. Geographic visualization was conducted using R (v4.4.3)^80^ and the ggplot2 package^81^. The coordinates were plotted onto a base map of Africa using the rnaturalearth package (scale = “medium”), and species-specific points were color-coded. All points were projected in the WGS84 coordinate reference system (EPSG:4326).

### Chromosome reordering and orientation standardization of public assemblies

For *S. indicum* ‘Xiaohei’ and *S. radiatum*, we reordered and reoriented the NCBI assemblies to match the published chromosome lengths and pairwise synteny reported by Wang et al. (2023)^17^. We generated whole-genome alignments using MUMmer4 nucmer v4.0.0 with parameters −l 20 (minimum exact match length) and -c 65 (minimum cluster length) for *S. indicum* ‘Xiaohei’ (accession GCA_027475695.1, retrieved 2023-01-04) and *S. radiatum* “*S. schinzianum* ‘Gangguo’” (accession GCA_027475655.1, retrieved 2023-01-04), and inferred syntenic relationships from these alignments. For each source chromosome, renaming and strand assignment (+/–) were determined by (i) the target chromosome with the largest covered length and (ii) the majority orientation of collinear blocks, while ensuring consistency with the chromosome length ranking reported in the literature. Subsequently, for *S. radiatum* (accession GCA_040286145.1, retrieved 2024-06-25) and *S. latifolium* (accession GCA_040286105.1, retrieved 2024-06-25) published by Miao et al. (2024)^8^, we used the harmonized *S. radiatum* “*S. schinzianum* ‘Gangguo’” assembly as the reference and applied the same procedure. Chromosome order and orientation were finalized based on one-to-one syntenic correspondence with *S. radiatum* “*S. schinzianum* ‘Gangguo’”, determined by the highest alignment coverage and consistent collinear block orientation. A chromosome naming and orientation concordance table is provided in Supplementary Table 8, listing for each public chromosome (by GenBank accession) the relative orientation used in this study (”+” indicates same orientation as the public record; “−” indicates reverse-complement relative to the public record) and the standardized chromosome name assigned in our analyses. No nucleotide sequences were modified; only chromosome names and orientations were standardized.

## Supporting information

Supplementary infomation

## Acknowledgements

We thank Prof. K. Kobayashi (Univ. Toyama) for his pioneer work on *Sesamum*. Dr. Yu Sugihara (Kyoto University) for phylogenetics. We thank Rie Ryusui (Itoh Laboratory, Institute of Science Tokyo) for her technical assistance with library preparation. This work was supported by JSPS KAKENHI Grant Number JP18K05571, JP16H06279 (PAGS), 22H02598.

