## Supplementary infomation for "An evolutionary landscape of sesame: chromosomal variation, allopolyploid speciation and metabolic specialization"

**Supplementary Information**

This Supplementary Information PDF includes Supplementary Methods, Supplementary Text, Supplementary Notes, Supplementary Figures 1–17, and Supplementary Tables 1, 2, 5, and 6. Owing to their size, Supplementary Tables 3, 4, 7, and 8 are provided as separate Excel files.

**Contents**

Supplementary Methods

Supplementary Text 1–2

Supplementary Figures 1–17

Supplementary Tables 1, 2, 5, and 6

### Supplementary Methods

#### LTR retrotransposon identification and phylogenetic analysis

Species-specific repeat libraries were generated for each genome assembly using RepeatModeler v2.0[1] with the -LTRStruct option enabled. Genome-wide repeat annotation was then carried out using RepeatMasker v4.0.7[2] with the de novo repeat library (-lib), soft masking (-xsmall), and GFF and alignment output options (-gff -a). Based on these annotations, Class I/LTR candidates were extracted and classified using TEsorter v1.4.6 with the REXdb-plant database and the -st nucl option. Domain-complete LTR retrotransposons were defined as entries annotated in the TEsorter summary table as Order = LTR and Complete = yes. Amino acid sequences corresponding to reverse transcriptase (RT) domains were extracted from the ".out.cls.pep" output files. Ty1-RT sequences were used for Copia-type elements and Ty3-RT sequences were used for Gypsy-type elements. Exact duplicate amino acid sequences were removed before alignment. Multiple sequence alignment was performed using MAFFT v7.407 with the --localpair and --maxiterate 1000 options. Poorly aligned sites were trimmed with trimAl v1.4 using the -gappyout option. Phylogenetic trees were inferred using VeryFastTree v4.0.5 under the LG+G model with the -lg -gamma options and enhanced search settings (-spr 4 -mlacc 2). Branch support was assessed by non-parametric bootstrap analysis with 1,000 replicates (-boot 1000). Trees were visualized in iTOL v6 using midpoint rooting while preserving original branch lengths. Lineage identity was indicated by branch colouring.

#### Genome-wide profiling of centromeric repeat (Cen1), LTR retrotransposons, and gene density

In this study, Cen1 refers to a centromeric repeat sequence originally identified as a satellite repeat enriched in centromeric regions of *Sesamum* by Miao et al. (2024). The Cen1 reference sequence was aligned to each assembled genome using BLASTN. Genomic regions showing significant sequence similarity to the Cen1 sequence were retained, and the corresponding genomic coordinates were extracted. These regions are hereafter referred to as Cen1-enriched regions and represent centromere-associated domains inferred from repeat accumulation rather than functionally validated centromeres. For genome-wide profiling, each chromosome was partitioned into non-overlapping windows of 1Mb. For each window, Cen1 coverage was calculated as the proportion of bases covered by Cen1 BLAST hits relative to window length. Continuous genomic intervals showing elevated Cen1 coverage were interpreted as regions enriched in centromeric repeats. The genomic distribution of LTR retrotransposons was analyzed on the basis of RepeatMasker annotations classified by TEsorter. LTR/Copia and LTR/Gypsy elements were extracted separately. For each genomic window, the total length annotated as each LTR category was summed and normalized by the window size to obtain coverage values. No filtering by insertion age was applied in these analyses. Gene density was calculated for the same set of windows using the final protein-coding gene annotation set. Genome-wide profiles of Cen1 repeat coverage, LTR/Copia coverage, LTR/Gypsy coverage, and gene density were visualized by concatenating chromosomes in a fixed order along the x-axis, enabling direct comparison of centromere-associated repeat accumulation, local transposable element composition, and gene distribution across the genome.

**Supplementary Text 1. Transposable element landscape across sesame relatives**

To compare transposable element (TE) composition among sesame relatives, we analyzed the genome assemblies of six newly sequenced species together with three previously reported references (*S. indicum* 'Xiaohei' and subgenomes A and B of *S. radiatum* "*S. schinzianum* 'Gangguo'"). Species-specific repeat libraries were first generated using RepeatModeler, and genome-wide repeat annotation was then performed with RepeatMasker, as described in Supplementary Methods. Across all genomes examined, LTR retrotransposons represented the predominant TE class (Supplementary Fig. 5; Supplementary Table 4). Among the sampled species, *S. alatum* had the highest LTR content (185.82 Mb) and also the largest genome size. By comparison, LTR content was lower in *C. triloba* (86.59 Mb), *S. latifolium* (90.73 Mb), and *S. indicum* 'Xiaohei' (38.87 Mb). Notably, the total size and composition of LTRs in *C. sesamoides* (61.28 Mb) and *S. angustifolium* (43.70 Mb) were similar to those of subgenomes A and B of *S. radiatum* "*S. schinzianum* 'Gangguo'" (60.64 Mb and 42.38 Mb, respectively). These results indicate broad correspondence in repeat composition between the inferred diploid relatives and the corresponding subgenomes of *S. radiatum*.

**Supplementary Text 2. Angela-type LTR/Copia expansion and Cen1-like repeat patterns in the DD lineage**

To investigate lineage-specific repeat dynamics, LTR retrotransposons were classified using TEsorter and their divergence profiles were compared among genomes. *S. alatum* showed a relative enrichment of Angela-type LTR/Copia elements, whereas Tekay/Athila-type LTR/Gypsy elements were comparatively moderate (Supplementary Figs. 15 and 16). Pairwise-divergence profiles further indicated a relatively young insertion peak for Angela-type elements in *S. alatum*, consistent with recent lineage-specific expansion (Supplementary Fig. 15). To examine chromosome-scale repeat organization, we also profiled the genomic distribution of Cen1-like repeats, LTR/Copia elements, LTR/Gypsy elements, and gene density across each genome using fixed genomic windows. Angela-type LTRs tended to accumulate in regions associated with Cen1-like repeats, and this relationship appeared altered in *S. alatum*, where extensive LTR proliferation coincided with reduced Cen1-associated signals (Supplementary Fig. 17). Together, these observations indicate that the DD lineage, represented by *S. alatum*, is characterized by a distinct repeat landscape with elevated LTR content and relative enrichment of Angela-type LTR/Copia elements. These genomic features coexist with the distinctive morphological and chemical characteristics reported for this lineage, including narrow leaves, a long floral tube, winged seeds, and specialized lignan chemotypes, although no direct causal relationship is inferred[3,4,5].

(A)

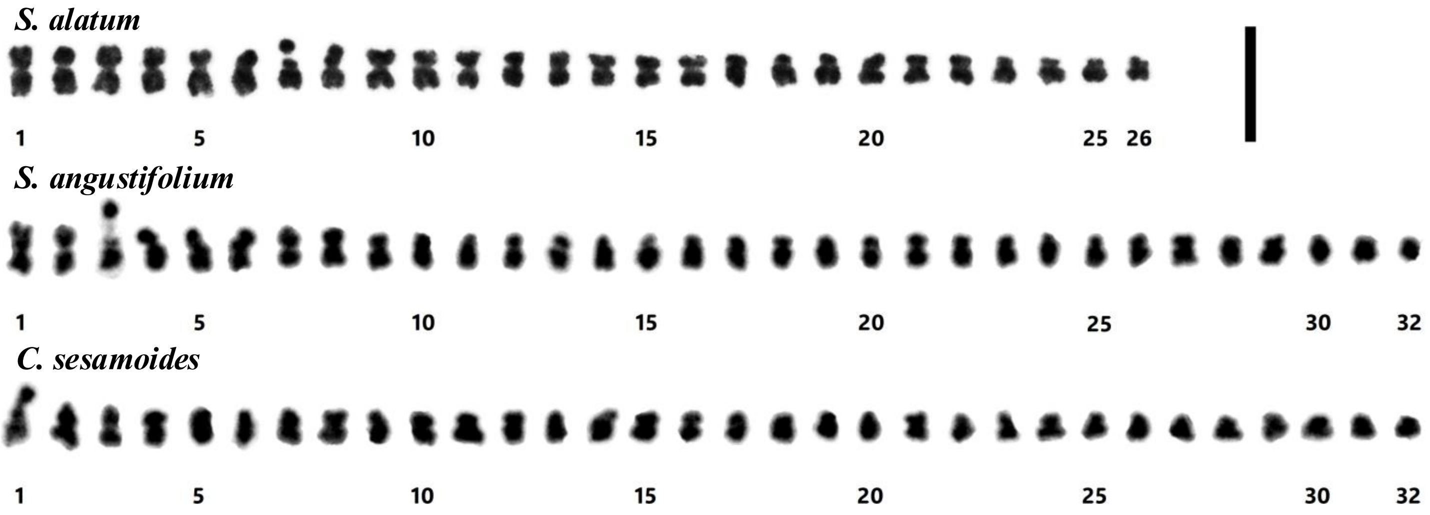

(B)

*S. alatum*

| No. | Length (μm) | Total (μm) | A.R. | Form |
|---|---|---|---|---|
| 1 | 0.9+1.1 | 2.0 | 1.2 | m |
| 2 | 0.9+1.1 | 2.0 | 1.2 | m |
| 3 | 0.9+1.0 | 1.9 | 1.1 | m |
| 4 | 0.9+1.0 | 1.9 | 1.1 | m |
| 5 | 0.8+1.1 | 1.9 | 1.4 | m |
| 6 | 0.8+1.1 | 1.9 | 1.4 | m |
| 7 | *0.6-0.3+1.0 | 1.9 | 1.1 | m |
| 8 | *0.6-0.3+1.0 | 1.9 | 1.1 | m |
| 9 | 0.8+0.9 | 1.7 | 1.1 | m |
| 10 | 0.8+0.9 | 1.7 | 1.1 | m |
| 11 | 0.8+0.9 | 1.7 | 1.1 | m |
| 12 | 0.8+0.9 | 1.7 | 1.1 | m |
| 13 | 0.7+0.8 | 1.5 | 1.1 | m |
| 14 | 0.7+0.8 | 1.5 | 1.1 | m |
| 15 | 0.6+0.7 | 1.3 | 1.2 | m |
| 16 | 0.6+0.7 | 1.3 | 1.2 | m |
| 17 | 0.5+0.8 | 1.3 | 1.6 | m |
| 18 | 0.5+0.8 | 1.3 | 1.6 | m |
| 19 | 0.5+0.8 | 1.3 | 1.6 | m |
| 20 | 0.5+0.8 | 1.3 | 1.6 | m |
| 21 | 0.5+0.6 | 1.1 | 1.2 | m |
| 22 | 0.5+0.6 | 1.1 | 1.2 | m |
| 23 | 0.4+0.6 | 1.0 | 1.5 | m |
| 24 | 0.4+0.6 | 1.0 | 1.5 | m |
| 25 | 0.4+0.5 | 0.9 | 1.3 | m |
| 26 | 0.4+0.5 | 0.9 | 1.3 | m |

*: Arms with satellites.

*S. angustifolium*

| No. | Length (μm) | Total (μm) | A.R. | Form |
|---|---|---|---|---|
| 1 | *0.2-0.7+1.0 | 1.9 | 1.1 | m |
| 2 | *0.2-0.7+1.0 | 1.9 | 1.1 | m |
| 3 | *0.6-0.3+1.0 | 1.9 | 1.1 | m |
| 4 | *0.6-0.3+1.0 | 1.9 | 1.1 | m |
| 5 | *0.6-0.3+1.0 | 1.9 | 1.1 | m |
| 6 | *0.6-0.3+1.0 | 1.9 | 1.1 | m |
| 7 | *0.2-0.6+0.9 | 1.7 | 1.1 | m |
| 8 | *0.2-0.6+0.9 | 1.7 | 1.1 | m |
| 9 | 0.6+0.8 | 1.4 | 1.3 | m |
| 10 | 0.6+0.8 | 1.4 | 1.3 | m |
| 11 | 0.5+0.8 | 1.3 | 1.6 | m |
| 12 | 0.5+0.8 | 1.3 | 1.6 | m |
| 13 | 0.5+0.8 | 1.3 | 1.6 | m |
| 14 | 0.5+0.8 | 1.3 | 1.6 | m |
| 15 | 0.4+0.8 | 1.2 | 2.0 | sm |
| 16 | 0.4+0.8 | 1.2 | 2.0 | sm |
| 17 | 0.4+0.7 | 1.1 | 1.8 | sm |
| 18 | 0.4+0.7 | 1.1 | 1.8 | sm |
| 19 | 0.4+0.7 | 1.1 | 1.8 | sm |
| 20 | 0.4+0.7 | 1.1 | 1.8 | sm |
| 21 | 0.4+0.6 | 1.0 | 1.5 | m |
| 22 | 0.4+0.6 | 1.0 | 1.5 | m |
| 23 | 0.3+0.7 | 1.0 | 2.3 | sm |
| 24 | 0.3+0.7 | 1.0 | 2.3 | sm |
| 25 | 0.3+0.6 | 0.9 | 2.0 | sm |
| 26 | 0.3+0.6 | 0.9 | 2.0 | sm |
| 27 | 0.3+0.6 | 0.9 | 2.0 | sm |
| 28 | 0.3+0.6 | 0.9 | 2.0 | sm |
| 29 | 0.2+0.6 | 0.8 | 3.0 | sm |
| 30 | 0.2+0.6 | 0.8 | 3.0 | sm |
| 31 | 0.3+0.4 | 0.7 | 1.3 | m |
| 32 | 0.3+0.4 | 0.7 | 1.3 | m |

*: Arms with satellites.

*C. sesamoides*

| No. | Length (μm) | Total (μm) | A.R. | Form |
|---|---|---|---|---|
| 1 | *0.6-0.3+1.0 | 1.9 | 1.1 | m |
| 2 | *0.6-0.3+1.0 | 1.9 | 1.1 | m |
| 3 | 0.7+0.9 | 1.6 | 1.3 | m |
| 4 | 0.7+0.9 | 1.6 | 1.3 | m |
| 5 | 0.6+0.8 | 1.4 | 1.3 | m |
| 6 | 0.6+0.8 | 1.4 | 1.3 | m |
| 7 | 0.6+0.7 | 1.3 | 1.2 | m |
| 8 | 0.6+0.7 | 1.3 | 1.2 | m |
| 9 | 0.6+0.7 | 1.3 | 1.2 | m |
| 10 | 0.6+0.7 | 1.3 | 1.2 | m |
| 11 | 0.6+0.6 | 1.2 | 1.0 | M |
| 12 | 0.6+0.6 | 1.2 | 1.0 | M |
| 13 | 0.5+0.6 | 1.1 | 1.5 | m |
| 14 | 0.5+0.6 | 1.1 | 1.5 | m |
| 15 | 0.4+0.5 | 0.9 | 1.3 | m |
| 16 | 0.4+0.5 | 0.9 | 1.3 | m |
| 17 | 0.4+0.5 | 0.9 | 1.3 | m |
| 18 | 0.4+0.5 | 0.9 | 1.3 | m |
| 19 | 0.3+0.6 | 0.9 | 2.0 | sm |
| 20 | 0.3+0.6 | 0.9 | 2.0 | sm |
| 21 | 0.3+0.5 | 0.8 | 1.6 | m |
| 22 | 0.3+0.5 | 0.8 | 1.6 | m |
| 23 | 0.3+0.5 | 0.8 | 1.6 | m |
| 24 | 0.3+0.5 | 0.8 | 1.6 | m |
| 25 | 0.3+0.4 | 0.7 | 1.3 | m |
| 26 | 0.3+0.4 | 0.7 | 1.3 | m |
| 27 | 0.3+0.4 | 0.7 | 1.3 | m |
| 28 | 0.3+0.4 | 0.7 | 1.3 | m |
| 29 | 0.3+0.4 | 0.7 | 1.3 | m |
| 30 | 0.3+0.4 | 0.7 | 1.3 | m |
| 31 | 0.3+0.4 | 0.7 | 1.3 | m |
| 32 | 0.3+0.4 | 0.7 | 1.3 | m |

*: Arms with satellites.

**Supplementary Fig. 1. Karyotypes of somatic metaphase of *S. alatum, S, angustifolium and C. sesamoides.***
**(A)** Karyograms of *S. alatum, S. angustifolium and C. sesamoides.* Scale Bar = 5 μm. Photomicrographs of somatic metaphase chromosomes of the three species are shown in Fig1.
**(B)** Measurements of somatic metaphase chromosomes of the species.

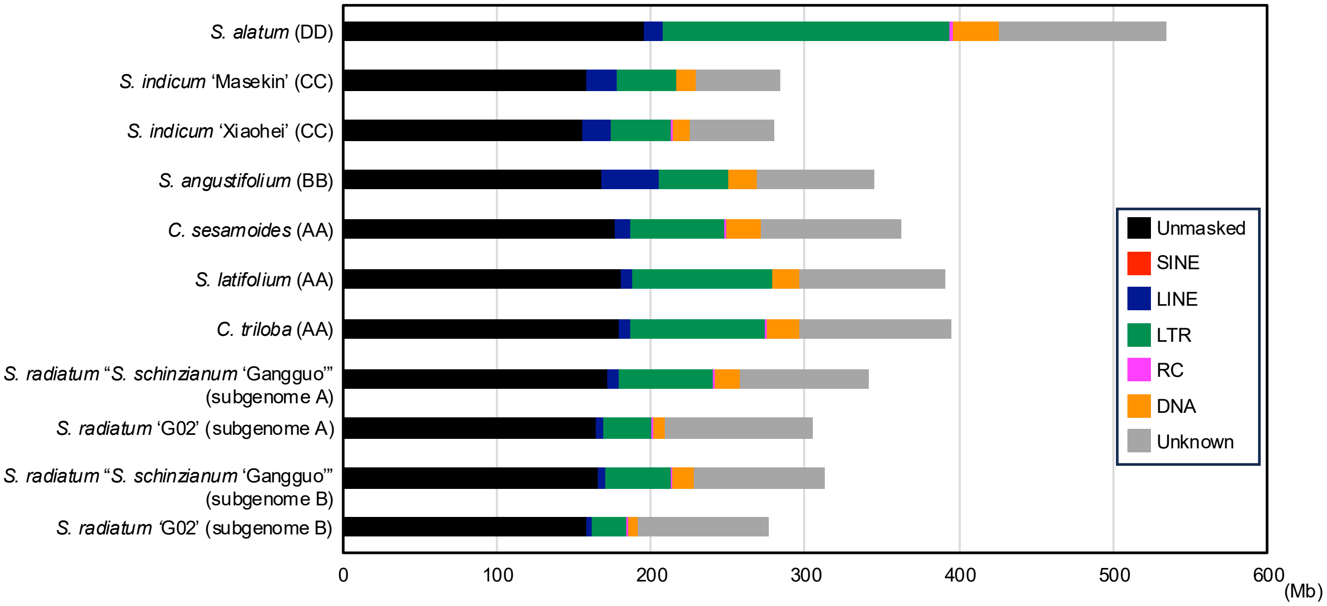

**Supplementary Fig. 2. Genome size and repeat composition across *Sesamum* and *Ceratotheca* species.** Each bar is partitioned by transposable-element (TE) category identified with RepeatMasker. Black segments denote unmasked (unique) sequence; coloured segments denote SINEs (red), LINEs (blue), LTR retrotransposons (green), Helitron/rolling-circle elements (magenta), DNA transposons (orange), and repeats of unknown classification (grey). The x-axis gives cumulative length in megabases (Mb).

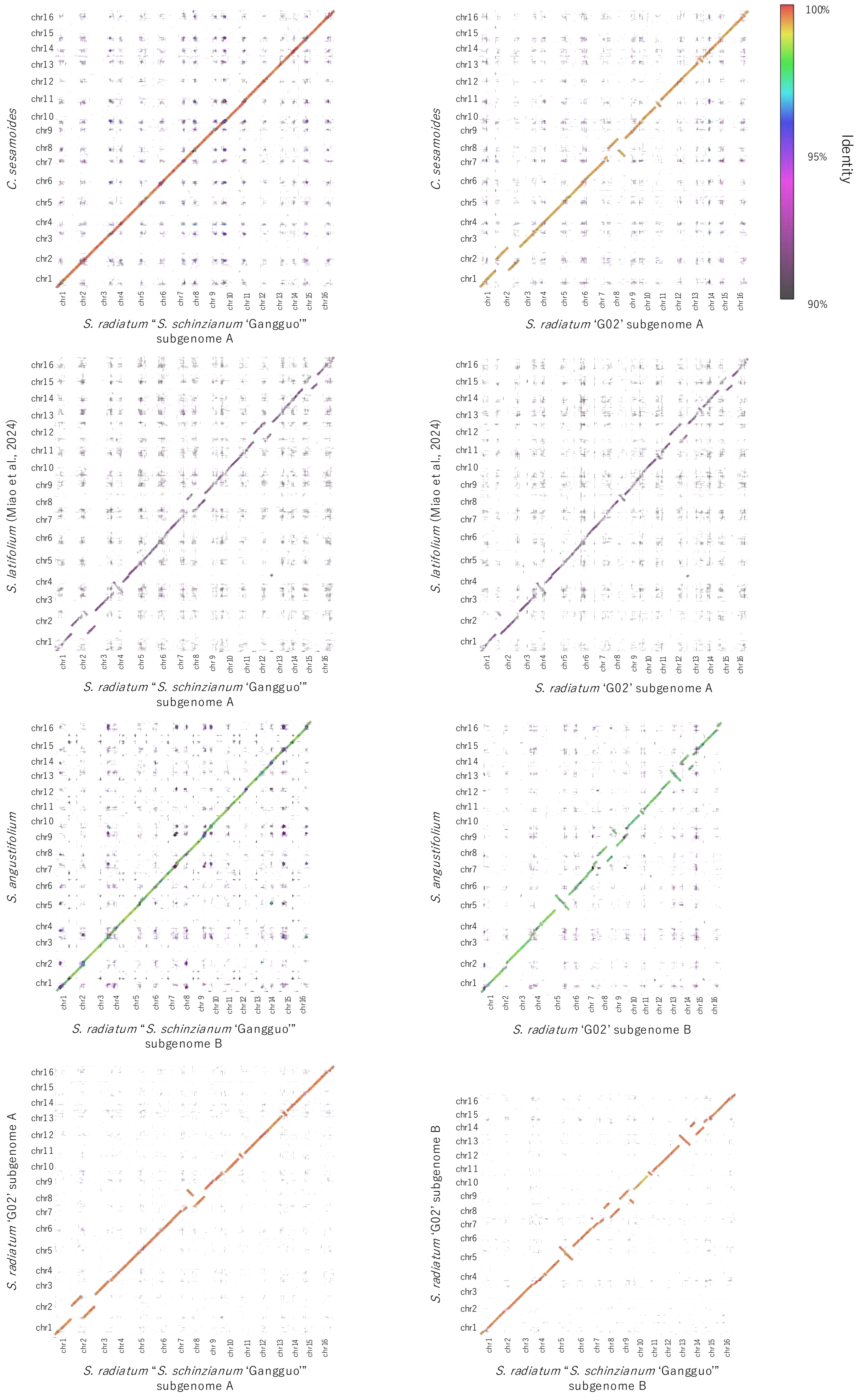

**Supplementary Fig. 3. Whole-genome dot plots showing nucleotide similarity between two species.** Each panel is a whole-genome dot plot for a pairwise comparison under the indicated setting. The x- and y-axes represent genomic coordinates of the concatenated chr of reference assembly and query assembly, respectively (scaffolds ordered by chromosome ID). Points mark local alignments retained at ≥ 1,000 bp computed with nucmer. Colors indicate alignment identity, as shown by the color bar. The genomes used for this analysis are as follows: *C. sesamoides* and *S. angustifolium* (this work), *S. radiatum* "*S. schinzianum* 'Gangguo'" (Wang et al., 2023), *S. latifolium*, and *S. radiatum* 'G02' (Miao et al., 2024).

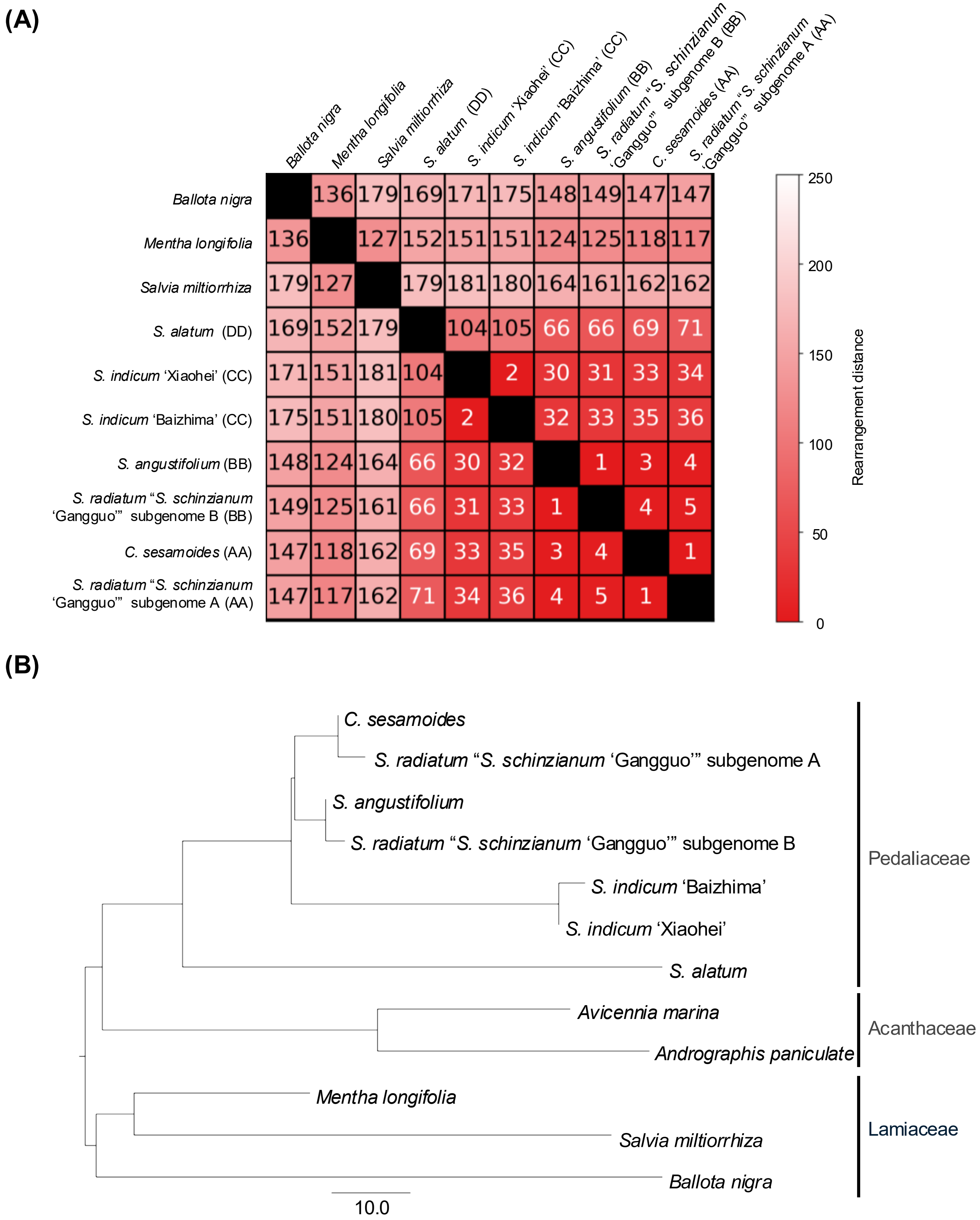

**Supplementary Fig. 4. Rearrangement-based phylogeny inferred using the CHROnicle framework.** Genome rearrangement analyses were performed using the CHROnicle pipeline based on genome-wide synteny blocks. (A) Pairwise rearrangement distance matrix estimated using ReChro. Values indicate the number of inferred chromosomal rearrangements separating each pair of genomes based on synteny block organization. Lower values (darker shading) denote higher structural similarity, whereas higher values reflect increased genome structural divergence. (B) Rearrangement-based phylogeny inferred using PhyChro, which reconstructs a species tree from the relative order and adjacency of conserved synteny blocks across genomes. The analysis includes 11 genomes from Pedaliaceae, Acanthaceae, and Lamiaceae, with subgenomes of Sesamum schinzianum treated separately. Taxonomic affiliations are indicated on the right.

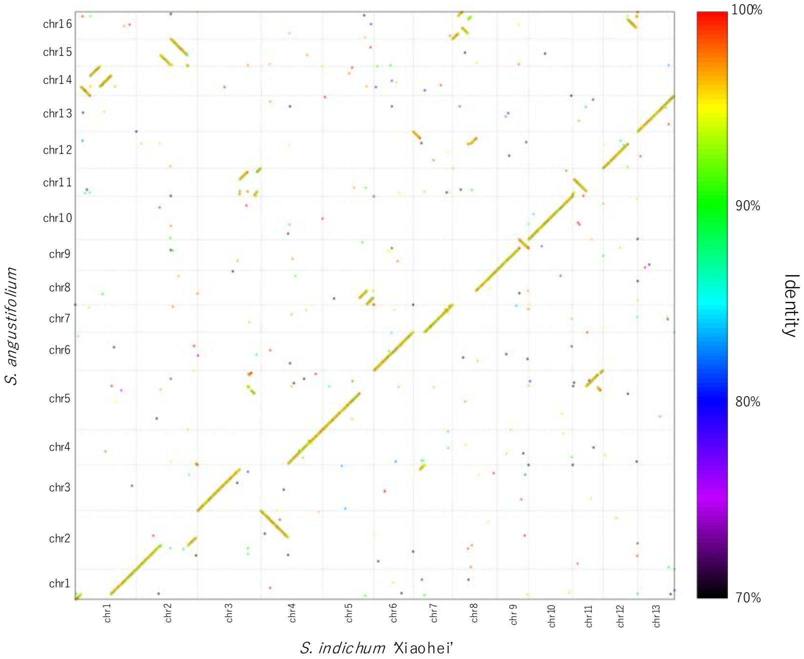

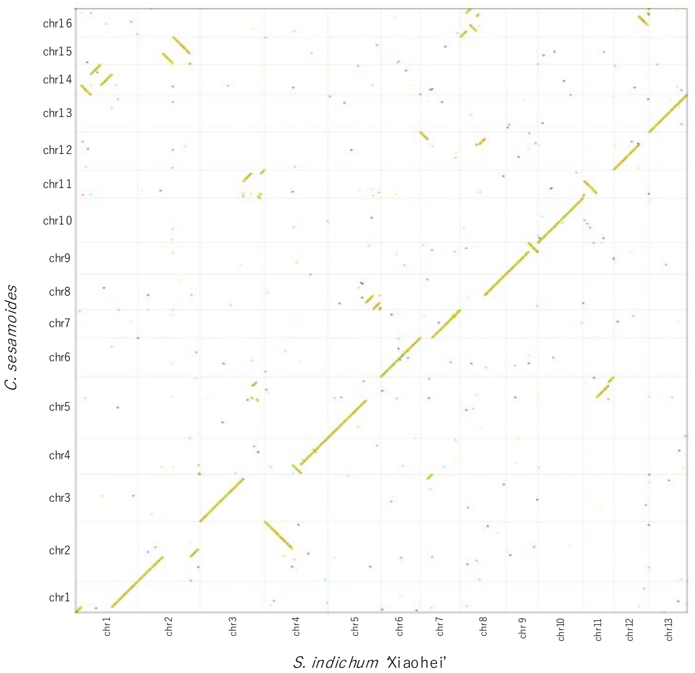

**Supplementary Fig. 5. Gene order conservation between *x* = 13 and *x* = 16 lineages.**
The upper panel shows the comparison between *S. angustifolium* and *S. indicum* 'Xiaohei' (20,815 reciprocal best-hit orthologous gene pairs), and the lower panel shows the comparison between *C. sesamoides* and *S. indicum* 'Xiaohei' (20,741 pairs). Each dot represents a reciprocal best-hit orthologous gene pair plotted according to cumulative gene order along pseudochromosomes. Dot colors indicate amino-acid sequence identity (%). Dotted grid lines indicate chromosome boundaries. Continuous diagonal patterns reflect conserved gene order, whereas interruptions and off-diagonal signals indicate chromosomal rearrangements and lineage-specific structural divergence.

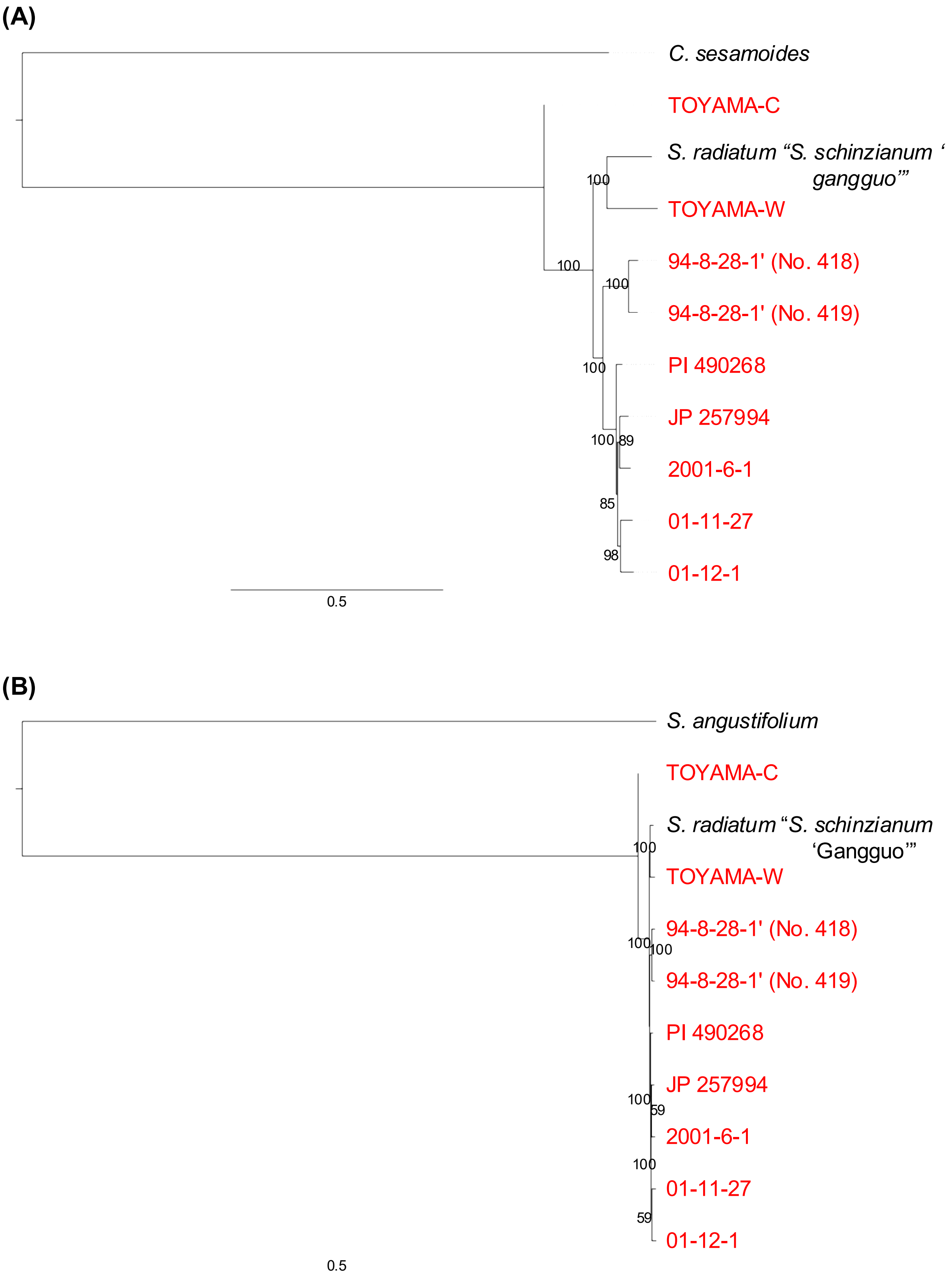

**Supplementary Fig. 6. Subgenome-specific phylogenies of *S. radiatum***

Maximum-likelihood phylogenetic trees were inferred from SNPs on (A) the A subgenome and (B) the B subgenome. Red labels indicate *S. radiatum* accessions. As putative progenitor lineages, *C. sesamoides* (A subgenome) and *S. angustifolium* (B subgenome) were included as outgroup. Numbers on the nodes indicate bootstrap value from 1,000 replications (left). The scale bar indicates the nucleotide substitutions per site.

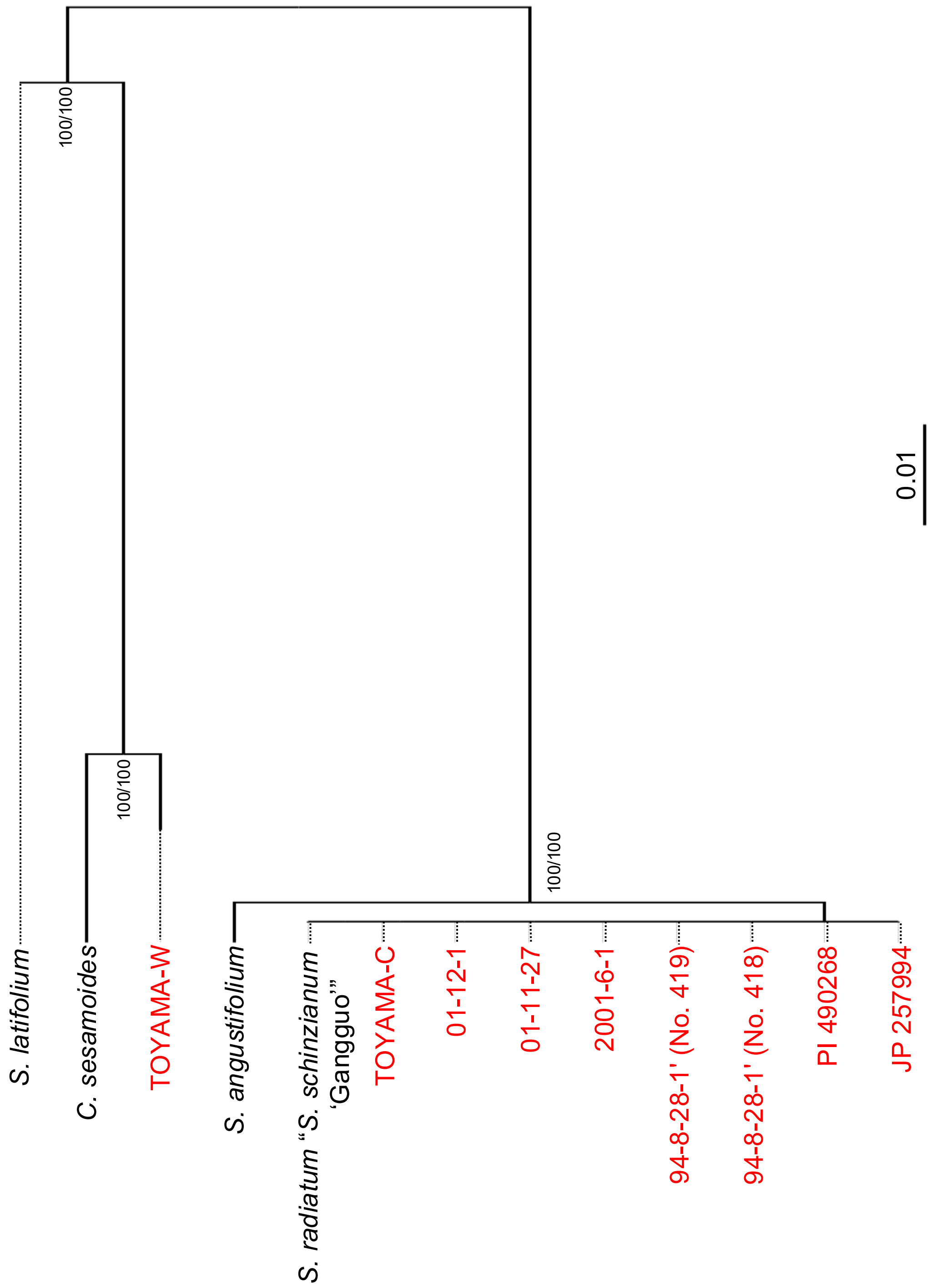

**Supplementary Fig. 7. Phylogenetic tree based on SNP positions in the mitochondria genome.** The line name of *S. radiatum* is highlighted in red. The scale bar indicates the nucleotide substitutions per site. Numbers on the nodes indicate bootstrap value from 1,000 replications (left), and SH-aLRT value (right). The scale bar indicates the substitutions per SNP site.

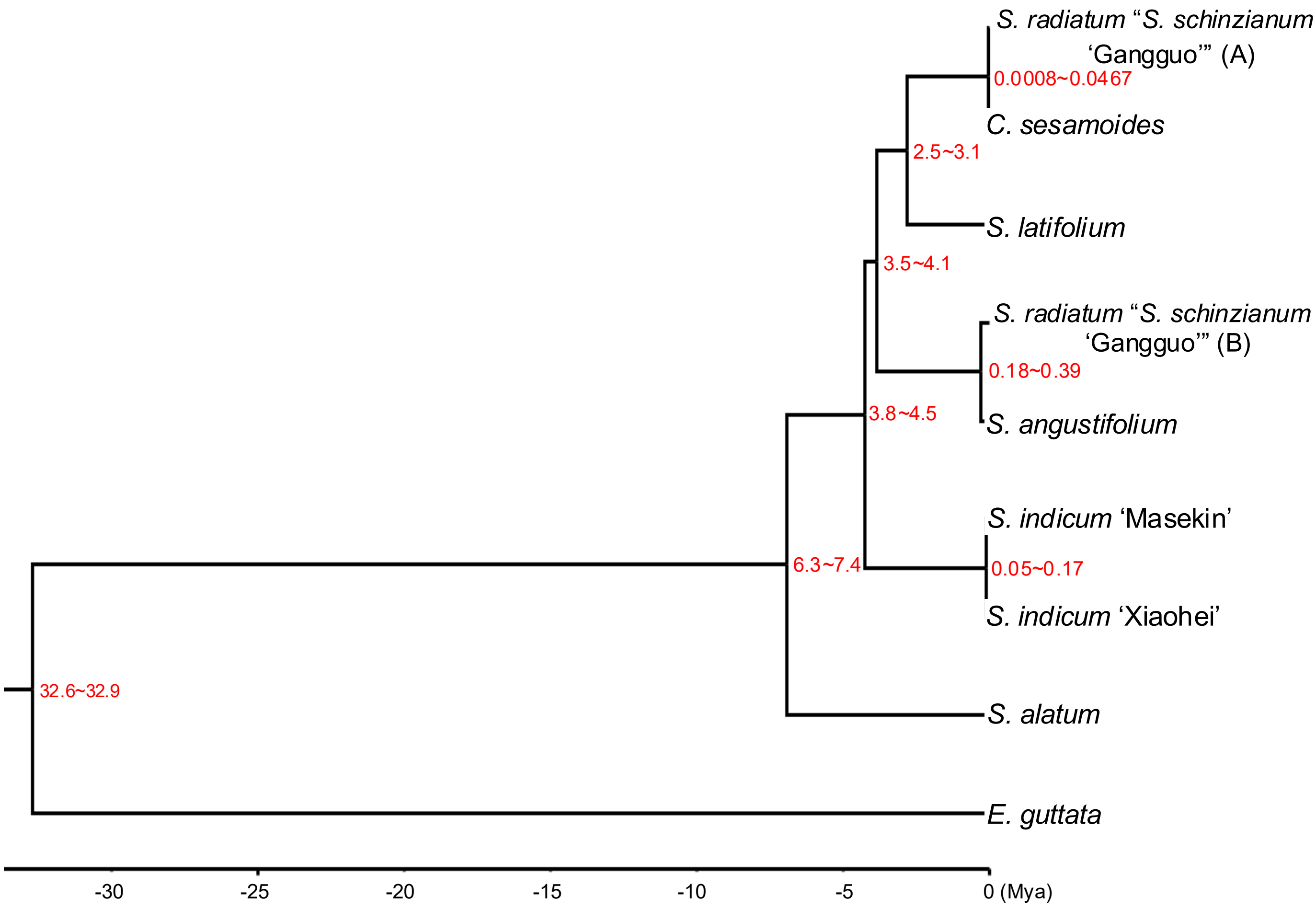

**Supplementary Fig. 8. Divergence-Time Estimation of *Sesamum* and *Ceratotheca*.**
Red node labels indicate the 95% highest posterior density (HPD) intervals for divergence times.

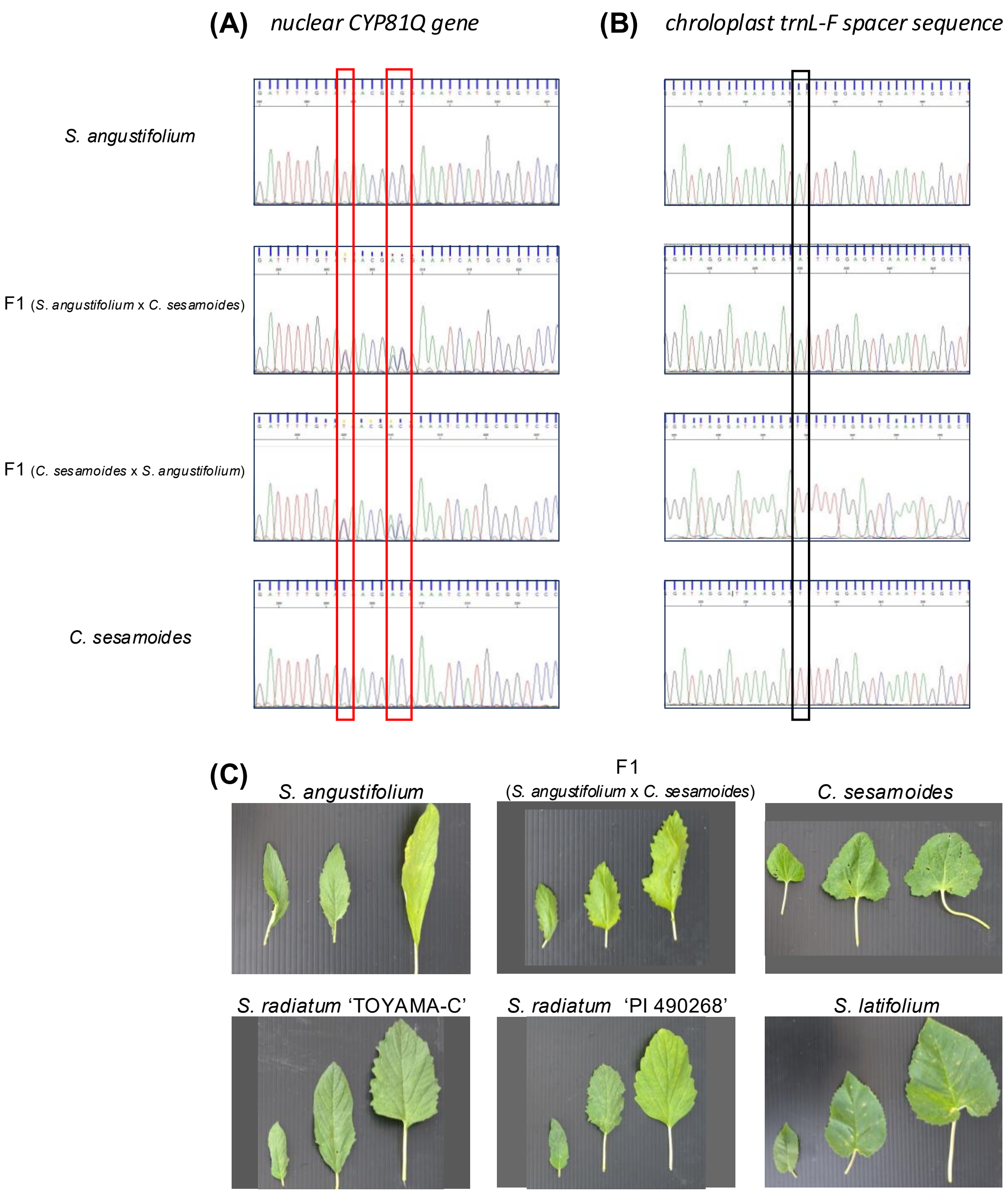

**Supplementary Fig. 9. Genetic and morphological characteristics of *S. angustifolium*, *C. sesamoides*, their F1 hybrid, and related species.**

(**A**) DNA sequence chromatograms of the CYP81Q1 region. The F1 hybrids are heterozygous, exhibiting double peaks at polymorphic sites where the parental species (*S. angustifolium* and *C. sesamoides*) are homozygous for different alleles. (**B**) DNA sequence chromatograms of the chloroplast trnL–trnF intergenic spacer. Species-diagnostic substitutions distinguish the parental species; the F1 hybrid carries a single chloroplast haplotype corresponding to the maternal parent. (**C**) Comparison of leaf morphology. The F1 hybrid (*S. angustifolium* x *C. sesamoides*) displays an intermediate morphology between the two parental species. Related species, *S. radiatum* (TOYAMA-C and PI 490268) and *S. latifolium*, are shown for comparison.

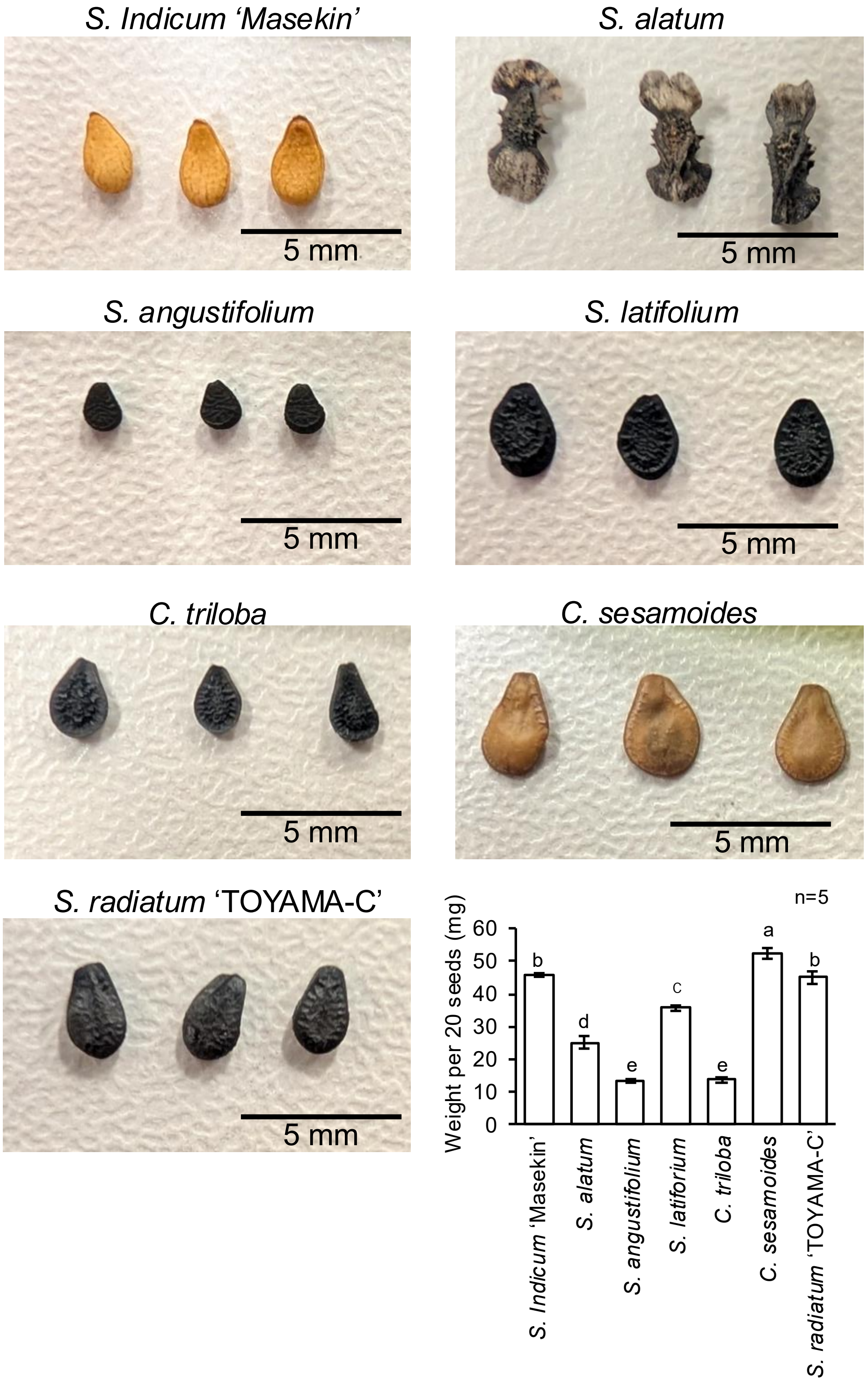

**Supplementary Fig. 10. Seed phenotypes of *Sesame*.**

Photographs show representative seeds of each species. *S. angustifolium* produces notably smaller seeds, whereas *C. sesamoides* exhibits a flatter seed morphology relative to the other species. The bar graph (lower right) shows the weight of 20 seeds; different letters denote significant differences among groups ($P < 0.05$, Tukey's HSD test).

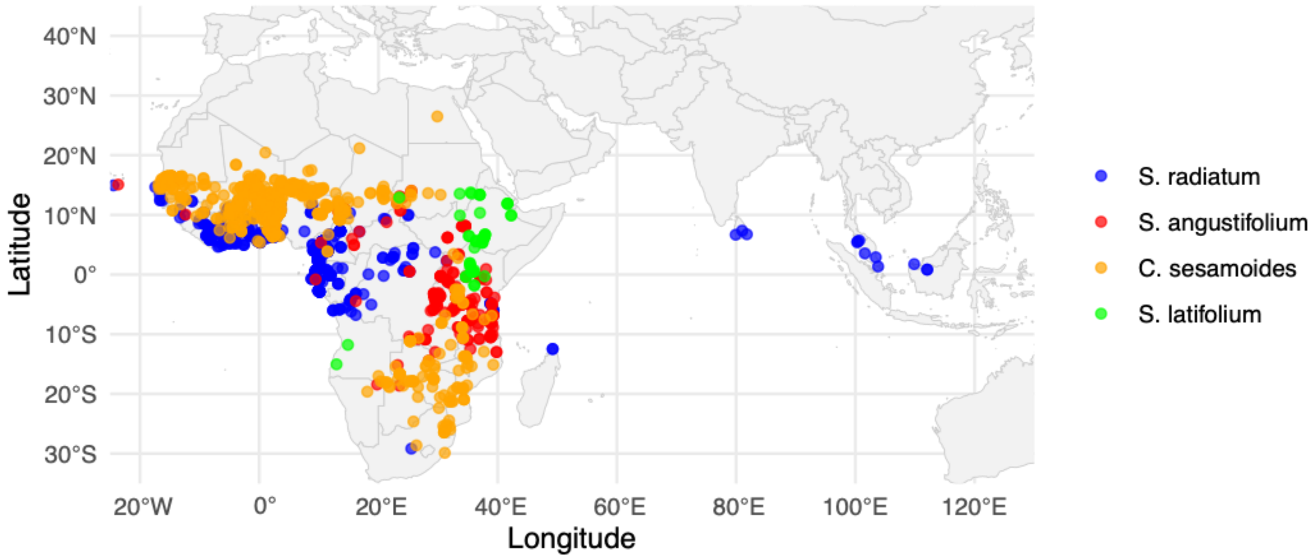

**Supplementary Fig. 11. Geographic distribution of *Sesamum* species and the related genus *Ceratotheca* in Africa and Southeast/East Asia.**

Occurrences of four taxa are plotted on a latitude–longitude grid (WGS-84, EPSG 4326). Each point marks a georeferenced herbarium or field record: orange, *C. sesamoides* (n = 2,253); red, *S. angustifolium* (n = 629); green, *S. latifolium* (n = 102); blue, *S. radiatum* (n = 1,310). Longitudinal lines are shown at 20° intervals from 20° W to 120° E; latitudinal lines at 10° intervals from 30° S to 40° N.

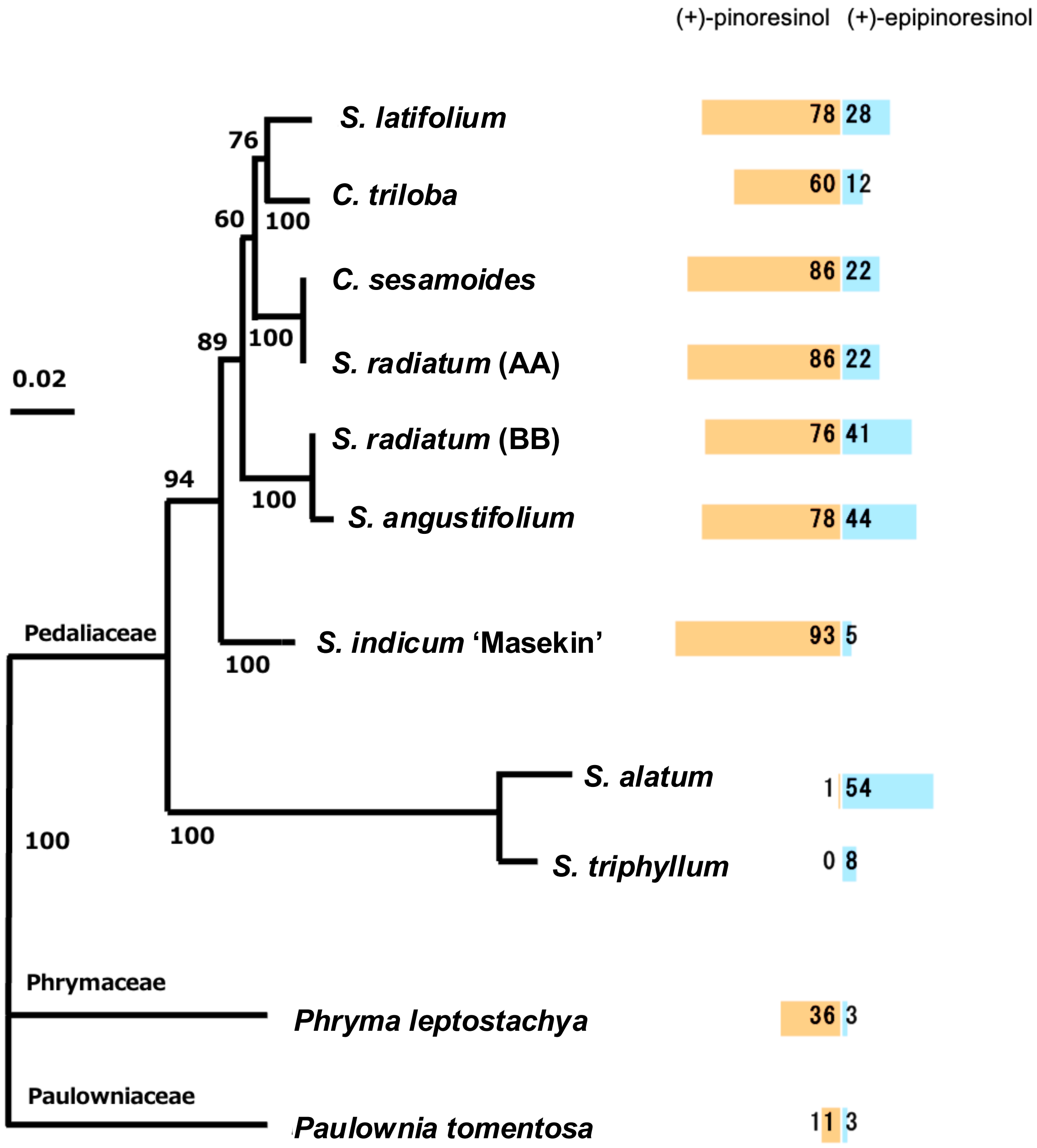

**Supplementary Fig. 12. Phylogeny of CYP81Q genes and their relative activities toward (+)-pinoresinol and (+)-epipinoresinol.**

The left panel depicts a maximum-likelihood tree reconstructed from concatenated single-copy nuclear genes. Numbers on branches are ultrafast-bootstrap support values ≥ 50 %; the scale bar corresponds to 0.02 expected amino-acid substitutions per site. Out-groups (Phryma leptostachya, Phrymaceae; Paulownia tomentosa, Paulowniaceae) are shown with dashed and green branches, respectively. Horizontal stacked bars on the right summarize in vitro catalytic activity of protein extracts toward the lignan precursors (+)-pinoresinol (tan) and (+)-epipinoresinol (light blue). Values indicate the mean specific activity (pmol product $min^{-1}$ $mg^{-1}$ protein; n = 3) attributed to each substrate; segments are scaled proportionally to the total activity measured for that species.

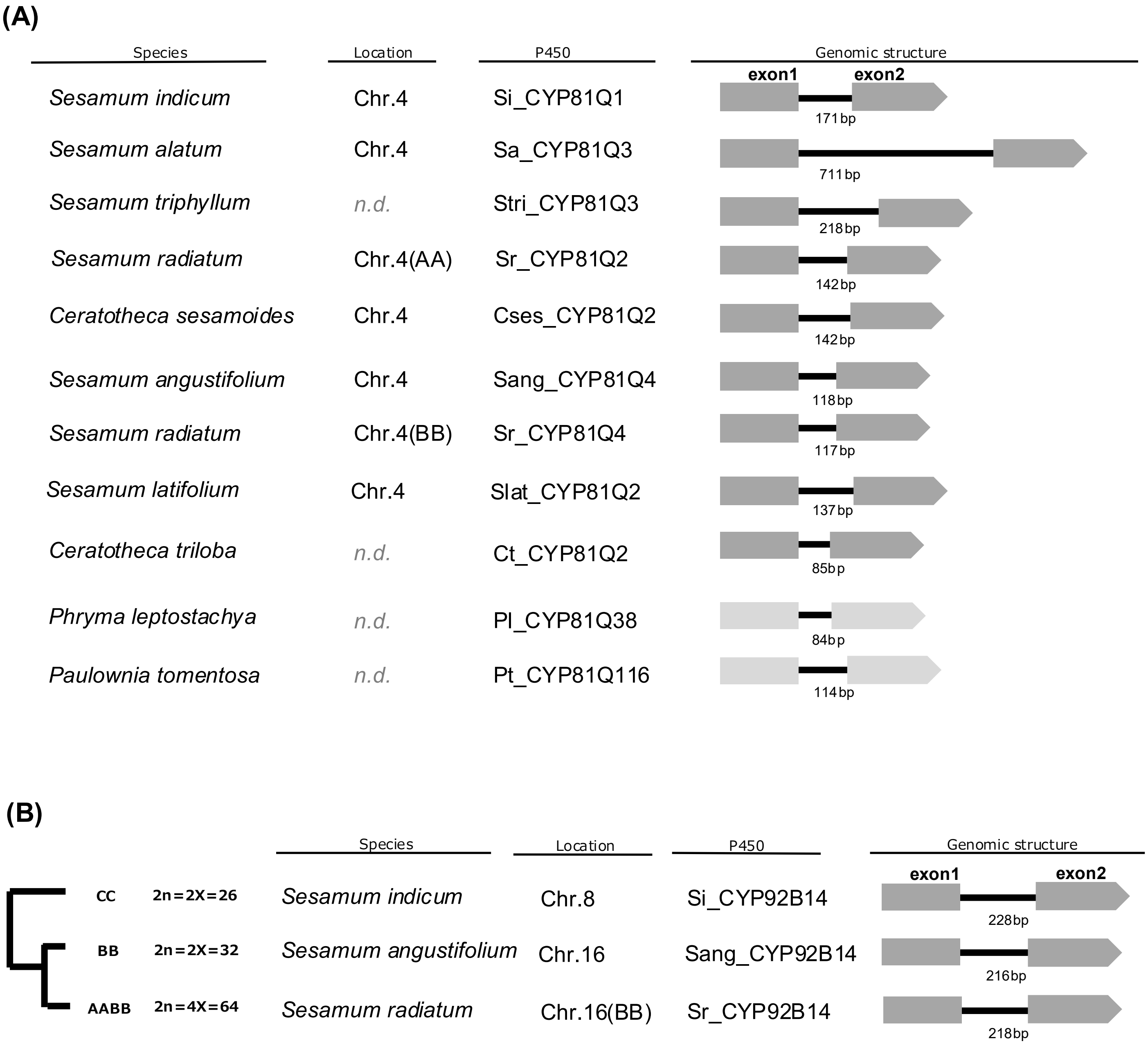

**Supplementary Fig. 13. Biosynthetic P450 genes for specialized sesame lignans**

(A) Summary of CYP81Q genes from *Sesamum* and its-related lignan producing plants of Lamiales.
All the these show a conserved structure of single intron between exon1 and exon2 and biochemical activity to form methylenedioxybridge on the substrate lignan.

(B) Summary of CYP92B14 genes in *Sesamum* plants which produce oxidized sesamin in their seeds.

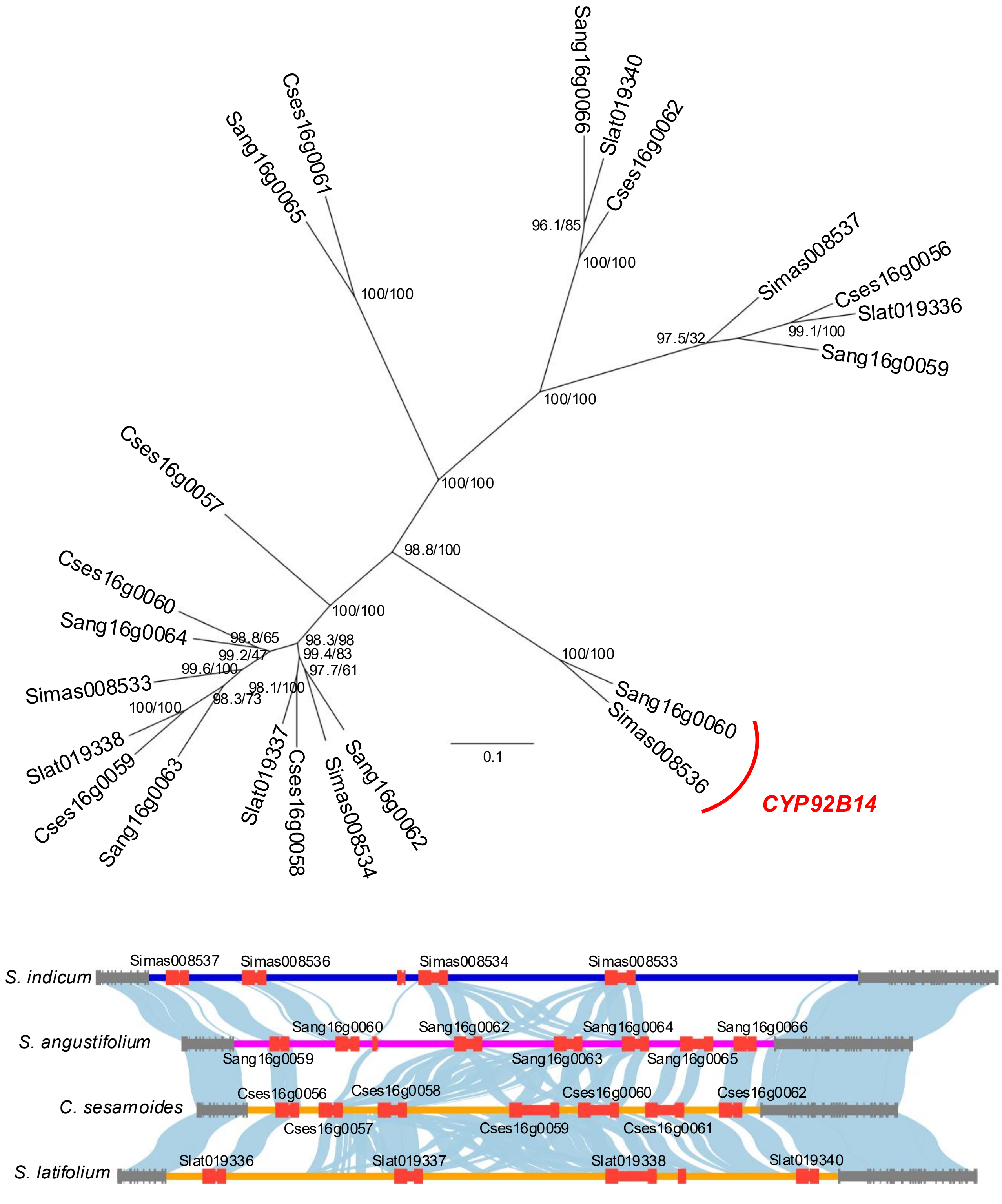

**Supplementary Fig. 14. Phylogeny of clustered P450 enzymes in the *CYP92B14* locus.**
The Maximum-likelihood tree was inferred from codon alignment sequences of P450s retrieved from the *CYP92B14* locus. Numbers on the nodes indicate bootstrap value from 1,000 replications (left), and SH-aLRT value (right). The scale bar indicates the substitutions per codon site. The branch labelled *CYP92B14* represents the functionally characterized (+)-sesamin oxidases.

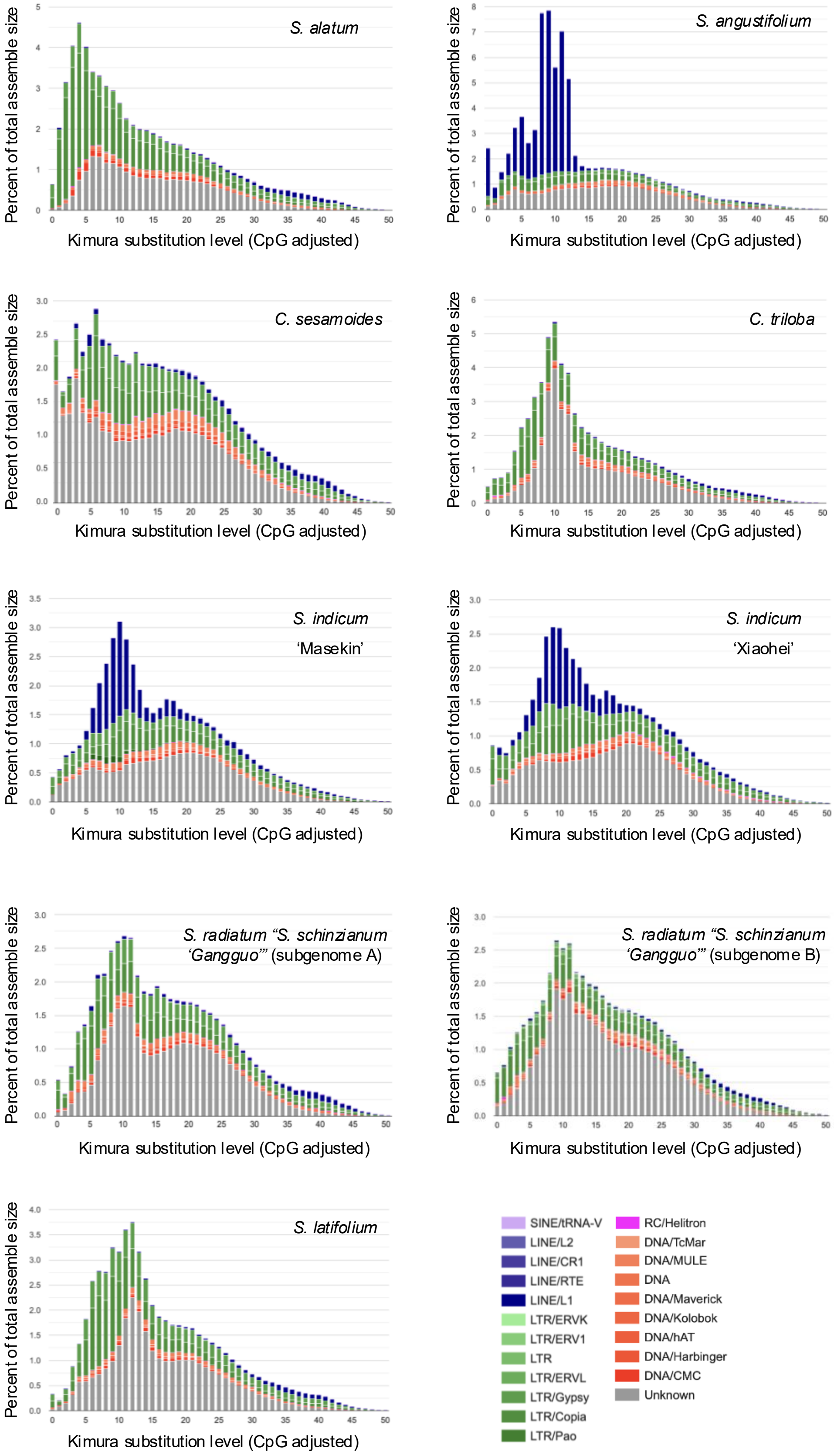

**Supplementary Fig. 15. Repeat landscapes of Pedaliaceae genomes.**
Stacked histograms show the genomic proportion of transposable-element (TE) classes across Kimura CpG-adjusted substitution levels (x-axis; one-per-cent bins). A higher Kimura's substitution level suggests an older evolutionary period. The y-axis indicates the percentage of the total assembly occupied by repeats in each bin.

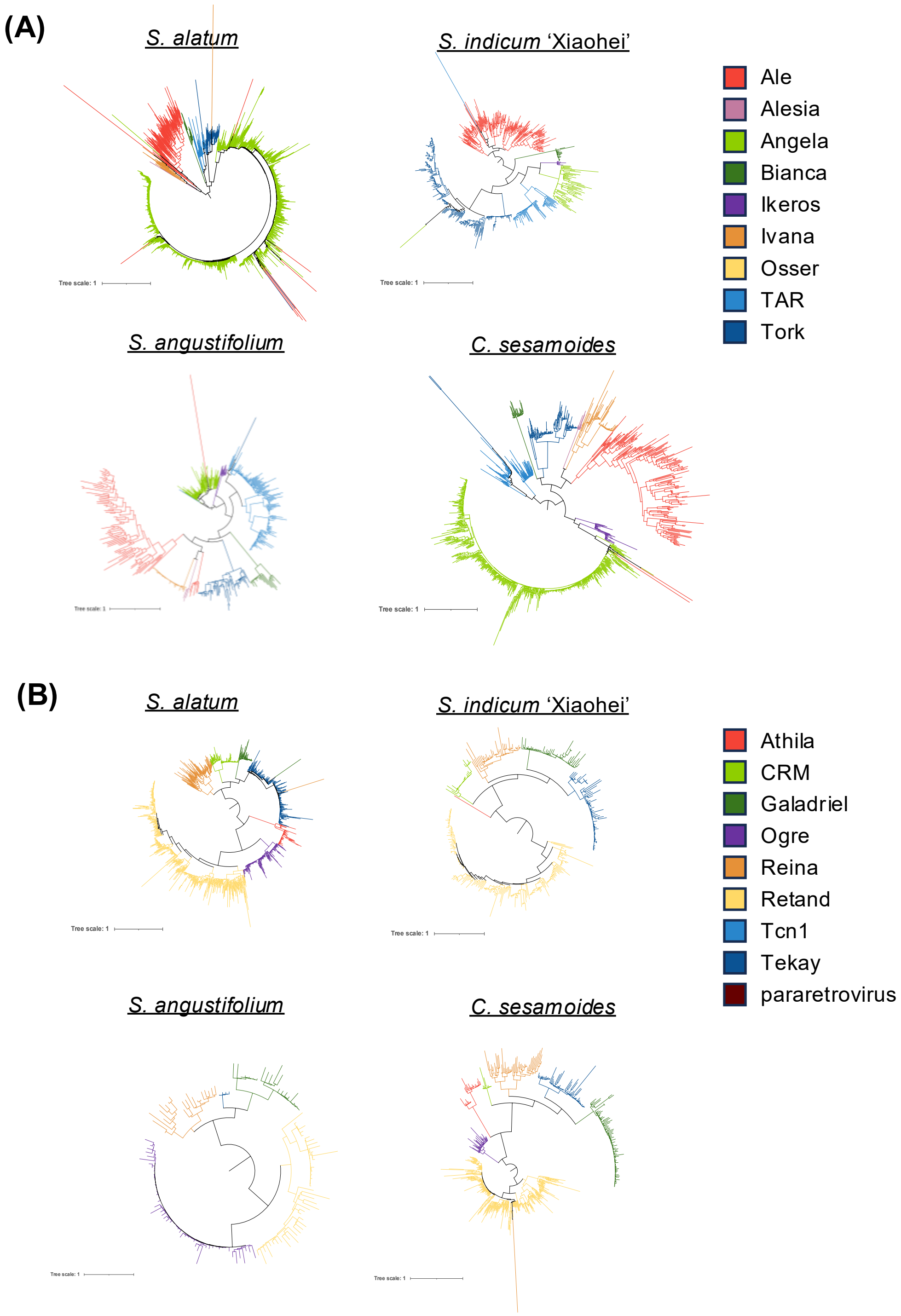

**Supplementary Fig. 16. Phylogenies of LTR retrotransposons based on RT domains.**
(A) Copia and (B) Gypsy lineages inferred from amino-acid alignments of the RT domain for two sesame genomes (*S. alatum* and *S. indicum* 'Xiaohei').

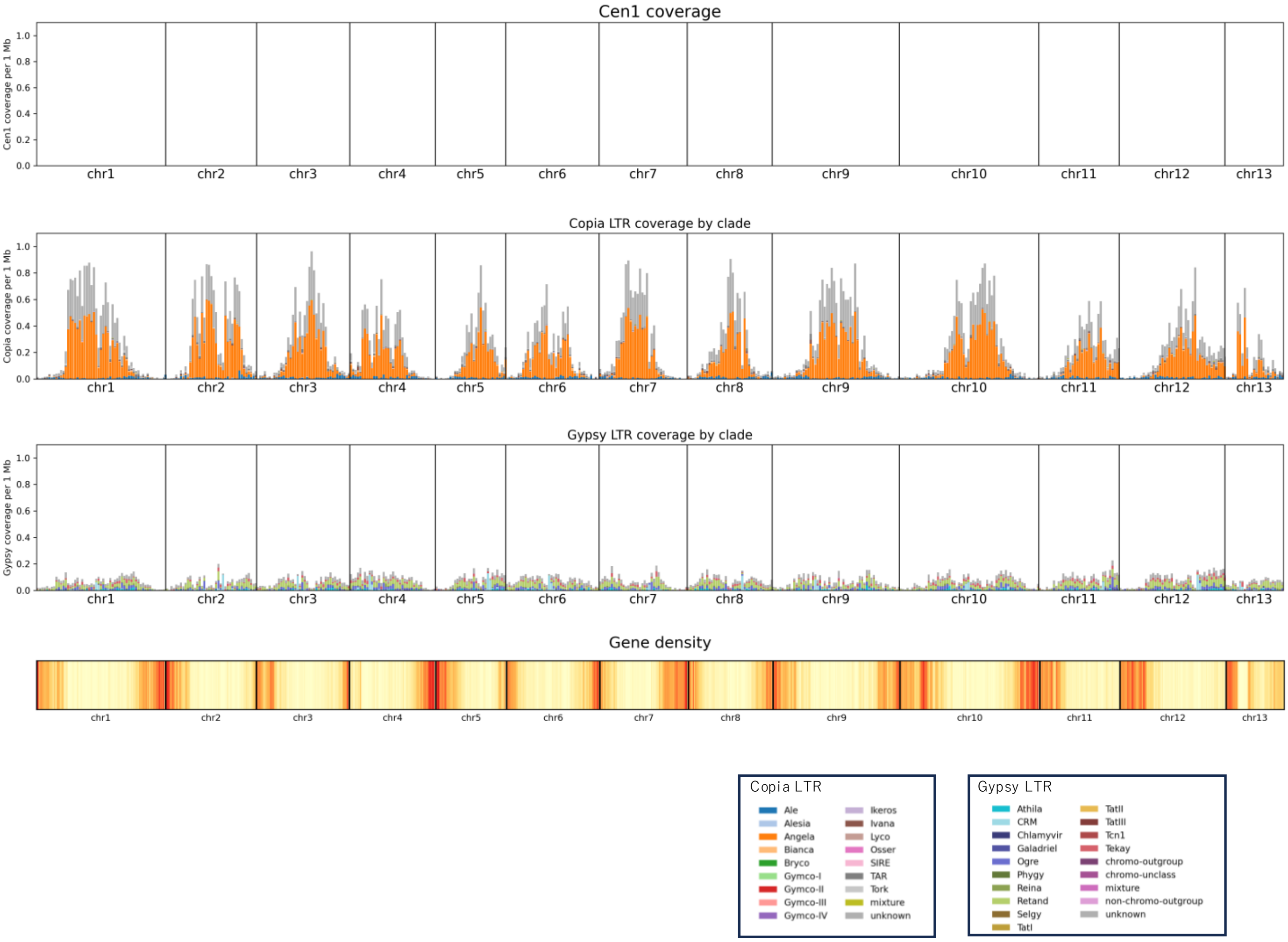

**Supplementary Fig. 17a. Distribution of Cen1-like repeats, LTR retrotransposons, and gene density across all chromosomes of *S. alatum***

Chromosome-scale distributions of Cen1-like centromeric repeat coverage, LTR/Copia and LTR/Gypsy coverage by clade, and gene density along all chromosomes (chr1–chr13) of *S. alatum*, calculated in non-overlapping 1 Mb windows.

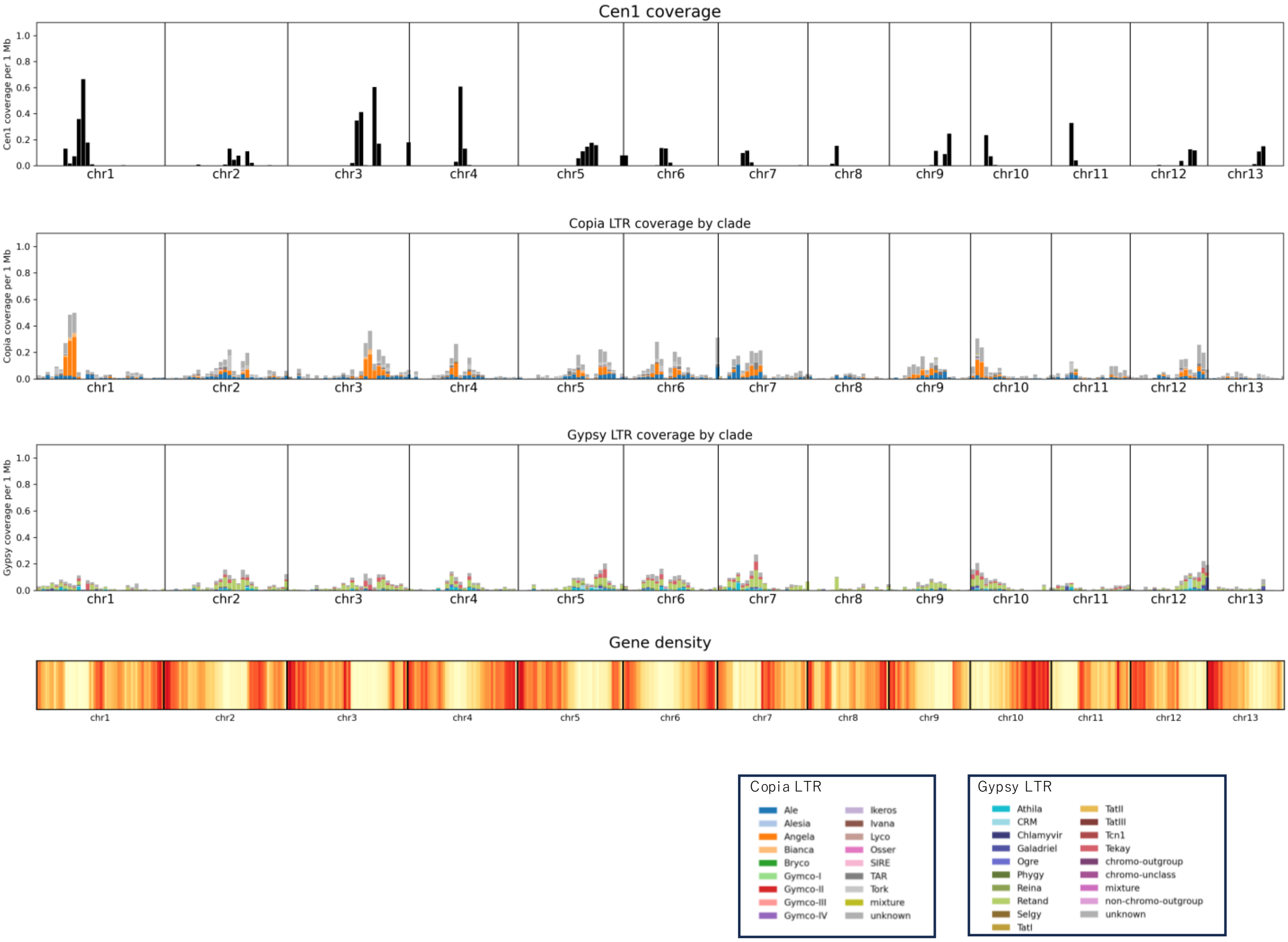

**Supplementary Fig. 17b. Distribution of Cen1-like repeats, LTR retrotransposons, and gene density across all chromosomes of *S. indicum* cv. Xiaohei**

Chromosome-scale distributions of Cen1-like centromeric repeat coverage, LTR/Copia and LTR/Gypsy coverage by clade, and gene density along all chromosomes (chr1–chr13) of *S. indicum* cv. Xiaohei, calculated in non-overlapping 1 Mb windows.

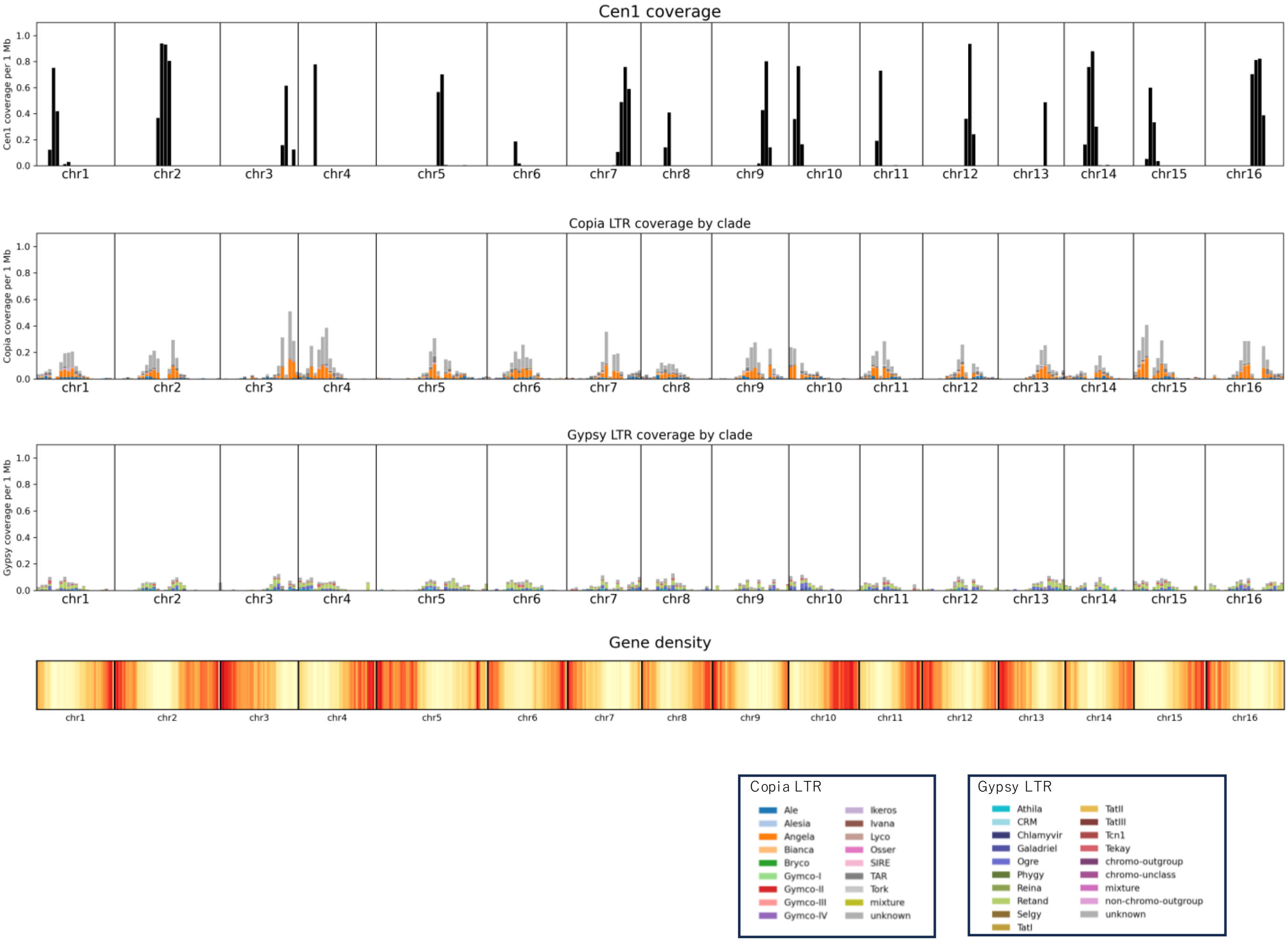

**Supplementary Fig. 17c. Distribution of Cen1-like repeats, LTR retrotransposons, and gene density across all chromosomes of *S. angustifolium***

Chromosome-scale distributions of Cen1-like centromeric repeat coverage, LTR/Copia and LTR/Gypsy coverage by clade, and gene density along all chromosomes (chr1–chr16) of *S. angustifolium*, calculated in non-overlapping 1 Mb windows.

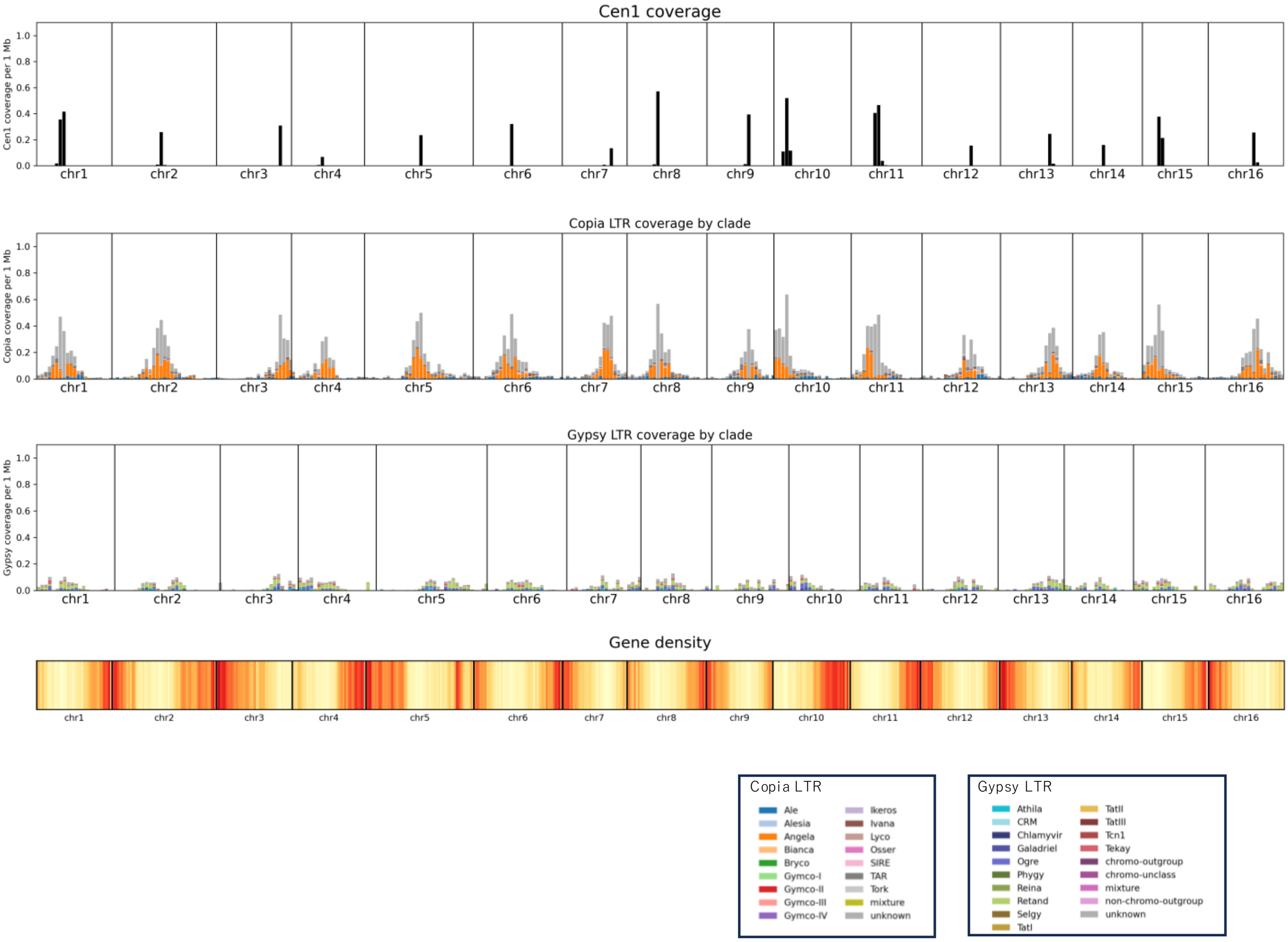

**Supplementary Fig. 17d. Distribution of Cen1-like repeats, LTR retrotransposons, and gene density across all chromosomes of *C. sesamoides***

Chromosome-scale distributions of Cen1-like centromeric repeat coverage, LTR/Copia and LTR/Gypsy coverage by clade, and gene density along all chromosomes (chr1–chr16) of *C. sesamoides*, calculated in non-overlapping 1 Mb windows.

**Supplementary Table 1. Sesame species used in this study**

| Species | Cultivar/accession | Chromosome numbers | Genome size (Mbp)[a] |
|---|---|---|---|
| ***S. indicum*** | Masekin | $2n = 2x = 26$ | 342.26 |
| ***S. alatum*** | Toyama Ala1[b] | $2n = 2x = 26$ | 536.93 |
| ***S. angustifolium*** | PI 367899 (USDA)[c] | $2n = 2x = 32$ | 354.88 |
| ***S. latifolium*** | Toyama Lat1[b] | $2n = 2x = 32$ | 439.72 |
| ***C. sesamoides*** | COL/GHANA/1993/MAFF/GJ93/233 | $2n = 2x = 32$ | 367.41 |
| ***C. triloba*** | PI 365013 (USDA) | $2n = 2x = 32$ | 407.01 |
| ***S. radiatum*** | | $2n = 4x = 64$ | |
| | PI 490268 (USDA) | | - |
| | JP 257994 | | - |
| | 94-8-28-1' (No. 418) | | - |
| | 94-8-28-1' (No. 419) | | - |
| | MAS2001-6-1 | | - |
| | MK01-11-27 | | - |
| | MK01-12-1 | | - |
| | TOYAMA-C [b] | | - |
| | TOYAMA-W [b, d] | | - |

[a] Estimated based on the k-mer frequency distribution

[b]Accessions conserved at University of Toyama

[c] Previously classified as *S. radiatum*

[d] Previously classified as *S. shinzianum*

**Supplementary Table 2. Accession numbers for genomes used in the study**

| Species | Cultivar/accession | BioSample | Assembly name | Accession number |
|---|---|---|---|---|
| *S. alatum* | Toyama Ala1 | SAMD00884009 | S_alatum_hic_v1.0 | AP039922-AP039950 |
| *S. angustifolium* | PI 367899 (USDA) | SAMD00884010 | S_angustifolium_hic_v1.0 | AP039951-AP040082 |
| *C.sesamoides* | COL/GHANA/1993/MAFF/GJ93/233 | SAMD00884013 | C.sesamoides_hic_v1.0 | AP039754-AP039921 |
| *C. triloba* | PI 365013(USDA) | SAMD00884012 | C_triloba_canu_draft_v1.0 | BAAGPD010000001-BAAGPD010000228 |
| *S. latifolium* | Toyama Lat1 | SAMD00884011 | S_latiforium_nextdenovo_draft_v1.0 | BAAGPF010000001-BAAGPF010000090 |
| *S. indicum* | Masekin | SAMD00884014 | S_indicum_cv_Masekin_nextdenovo_draft_v1.0 | BAAGPE010000001-BAAGPE010000098 |

**Supplementary Table 5. Lignan contents in the seeds of sesame species**

| | 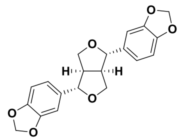 | 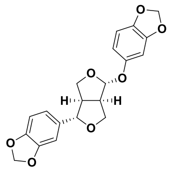 | 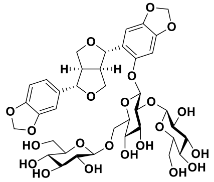 | 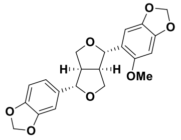 | 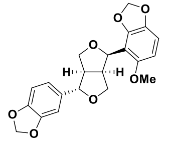 | 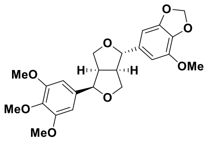 | 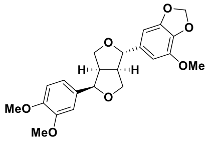 |
|---|---|---|---|---|---|---|---|
| | (+)-sesamin | (+)-sesamolin | SL-TG* | (+)-sesangolin | (+)-7'-episesantalin | (+)-2-episesalatin | (+)-alatumin |
| *S. indicum* | 5.34 | 2.93 | 0.57 | N.D. | N.D. | N.D. | N.D. |
| *S. alatum* | N.D. | N.D. | N.D. | N.D. | N.D. | 2.14 | 2.03 |
| *S. angustifolium* | 0.01 | 0.27 | N.D. | 8.47 | 2.36 | N.D. | N.D. |
| *S. latifolium* | 10.45 | N.D. | N.D. | N.D. | N.D. | N.D. | N.D. |
| *C. sesamoides* | 10.29 | N.D. | N.D. | N.D. | N.D. | N.D. | N.D. |
| *C. triloba* | N.D. | N.D. | N.D. | N.D. | N.D. | N.D. | N.D. |
| *S. radiatum* TOYAMA-C | 0.32 | 0.02 | N.D. | 2.38 | 0.14 | N.D. | N.D. |
| *S. radiatum* TOYAMA-W | 1.23 | 0.10 | N.D. | 5.46 | 0.30 | N.D. | N.D. |
| *S. radiatum* PI 490268 (USDA) | 1.73 | 0.10 | N.D. | 5.06 | 0.31 | N.D. | N.D. |
| *S. radiatum* JP 257994 | 1.09 | 0.08 | N.D. | 3.80 | 0.24 | N.D. | N.D. |
| *S. radiatum* 94-8-28-1' (No. 418) | 1.18 | 0.10 | N.D. | 4.49 | 0.26 | N.D. | N.D. |
| *S. radiatum* 94-8-28-1' (No. 419) | 1.54 | 0.11 | N.D. | 5.13 | 0.30 | N.D. | N.D. |

*SL-TG: (+)-sesaminol-triglucoside

N.D.: not detected

**Supplementary Table 6. Accession numbers for sequence data used in the study**

| Species | Cultivar/accession | BioSample | Accession number | Library types | Reads number | Raw data (bp) | Estimate read coverage |
|---|---|---|---|---|---|---|---|
| *S. alatum* | Toyama Ala1 | SAMD00884009 | DRR640264 | Illumina paired-end | 315,795,172 | 47,369,275,800 | 88.2 |
| | | | DRR640265 | PacBio CLR | 6,790,097 | 174,394,058,538 | 324.8 |
| | | | DRR640266 | Hi-C (Omni-C) | 163,314,114 | 24,660,431,214 | 45.9 |
| *S. angustifolium* | PI 367899 (USDA) | SAMD00884010 | DRR640267 | Illumina paired-end | 355,031,872 | 53,254,780,800 | 150.1 |
| | | | DRR640268 | PacBio CLR | 7,831,509 | 188,228,844,978 | 530.4 |
| | | | DRR640269 | Hi-C (Omni-C) | 173,089,584 | 26,136,527,184 | 73.6 |
| *C. sesamoides* | COL/GHANA/1993/MAFF/GJ93/233 | SAMD00884013 | DRR640270 | Illumina paired-end | 286,253,300 | 42,937,995,000 | 116.9 |
| | | | DRR640271 | PacBio CLR | 8,192,506 | 167,329,512,383 | 455.4 |
| | | | DRR640272 | Hi-C (Omni-C) | 154,693,114 | 23,358,660,214 | 63.6 |
| *C. triloba* | PI 365013(USDA) | SAMD00884012 | DRR640273 | Illumina-PE | 468,788,526 | 70,318,278,900 | 172.7 |
| | | | DRR640274 | PacBio CLR | 8,240,035 | 155,869,228,700 | 383 |
| *S. latifolium* | Toyama Lat1 | SAMD00884011 | DRR640275 | Illumina-PE | 856,613,934 | 128,492,090,100 | 292.2 |
| | | | DRR640276 | Nanopore | 7,139,381 | 52,402,441,485 | 119.2 |
| *S. indicum* | Masekin | SAMD00884014 | DRR640277 | Illumina-PE | 856,546,660 | 128,481,999,000 | 375.4 |
| | | | DRR640278 | Nanopore | 6,608,552 | 71,498,513,647 | 208.9 |
| *S. radiatum* | PI 490268 (USDA) | SAMD00884015 | DRR640279 | Illumina-PE | 83,798,420 | 12,569,763,000 | 18.8 |
| | JP 257994 | SAMD00884016 | DRR640280 | Illumina-PE | 88,217,264 | 13,232,589,600 | 19.8 |
| | 94-8-28-1' (No. 418) | SAMD00884017 | DRR640281 | Illumina-PE | 76,597,822 | 11,489,673,300 | 17.2 |
| | 94-8-28-1' (No. 419) | SAMD00884018 | DRR640282 | Illumina-PE | 53,225,418 | 7,983,812,700 | 11.9 |
| | MAS2001-6-1 | SAMD00884019 | DRR640283 | Illumina-PE | 49,561,130 | 7,434,169,500 | 11.1 |
| | MK01-11-27 | SAMD00884020 | DRR640284 | Illumina-PE | 85,223,208 | 12,783,481,200 | 19.1 |
| | MK01-12-1 | SAMD00884021 | DRR640285 | Illumina-PE | 118,789,980 | 17,818,497,000 | 26.7 |
| | TOYAMA-C | SAMD00884022 | DRR640286 | Illumina-PE | 28,100,538 | 4,215,080,700 | 6.3 |
| | TOYAMA-W | SAMD00884023 | DRR640287 | Illumina-PE | 86,815,234 | 13,022,285,100 | 19.5 |
